# ESMDynamic: Fast and Accurate Prediction of Protein Dynamic Contact Maps from Single Sequences

**DOI:** 10.1101/2025.08.20.671365

**Authors:** Diego E. Kleiman, Jiangyan Feng, Zhengyuan Xue, Diwakar Shukla

## Abstract

Understanding conformational dynamics is essential for elucidating protein function, yet most deep learning models in structural biology predict only static structures. Here, we present ESMDynamic, a deep learning model that predicts residue–residue contact dynamics directly from protein sequence. Built on the ESMFold architecture and trained on conformational variability from experimental structure ensembles and molecular dynamics (MD) simulations, ESMDynamic predicts dynamic contact probabilities, contact frequencies reflecting equilibrium populations, and coarse-grained kinetics of contact formation and dissociation across multiple temperature conditions. On large-scale MD benchmarks (mdCATH and ATLAS), ESMDynamic matches or outperforms state-of-the-art ensemble prediction methods (AlphaFlow, ESMFlow, BioEmu) while requiring orders-of-magnitude less computation. We demonstrate generalization to diverse systems, including membrane transporters, a *de novo* designed protein, and a homodimer complex. We show that predicted dynamic contacts enable automated selection of collective variables for Markov state model construction. Applied to the human proteome, ESMDynamic generates predictions for over 18,000 proteins, enabling large-scale analysis of conformational variability. Overall, ESMDynamic provides a scalable, sequence-based representation of protein dynamics to inform simulation, analysis, and design workflows.

## Introduction

In recent years, machine-learning (ML) models have achieved excellent accuracy in predicting protein structure from amino acid sequences by exploiting their encoded evolutionary information. While some models^1–5^ utilize multiple-sequence alignments (MSAs)^6,7^ to acquire this information, other models employ protein language models (PLMs)^8–10^ to embed sequences in a latent space whose features facilitate downstream tasks, including structure prediction. Proteins always exist as conformational ensembles in thermodynamic equilibrium.^11^ While knowledge of protein structure is crucial for understanding function, the current paradigm establishes the role of dynamics as a necessary hinge point between structure and function,^12^ a relationship that was determined as early as the first cooperative binding models^13^ and was later verified in many instances of allosteric regulation,^14^ motor proteins,^15^ and enzymatic catalysis,^16^ among others.^17–20^ However, structure prediction models only output static structures. Even in cases where a sequence maps to different stable conformations,^21^ these models tend to suffer from mode collapse,^22^ where a single high-likelihood prediction is favored over other possible outputs. Nonetheless, several methodological innovations made it possible to predict multiple structures from sequences.^23–27^

Due to the relevance of dynamics in the elucidation of protein function, many methods have been devised to predict structural ensembles. Many such protocols rely on partially hiding, masking, or noising the coevolutionary data passed to the models in a random^23^ or a biased^24^ fashion to introduce diversity among the predictions. However, these methods present certain shortcomings. Firstly, the model must be iteratively queried with the perturbed inputs, increasing the inference time. Secondly, the sensitivity of the models to input perturbations is poorly characterized. Therefore, it cannot be known *a priori* whether alternative conformations will be predicted, and if they are, it is challenging to design the perturbation that results in the desired output change. Some authors have suggested that this approach is successful only when the alternative conformations are observed during training,^22^ although results with held-out test sets indicate the contrary.^3^ These methods have been extended by incorporating molecular dynamics (MD) simulations to sample a free energy landscape of the system once a conformational ensemble has been predicted.^28^ Another family of methods employs diffusion-based or flow-based probabilistic models to sample conformational ensembles.^26,27^

A different approach to predict alternative conformations relies on reinterpreting the output of structure prediction models to infer the implicit conformational diversity encoded within intermediate outputs, which is typically suppressed once the model converges on a final structure. These approaches may construct energy restrictions based on pairwise residue distances.^29^ However, these approaches are only intended to work on capturing large conformational changes, mostly rearrangements of interdomain distances. Nonetheless, many functionally relevant transition are better described as subtle fluctuations propagated through networks of dynamically-coupled residues.^30^

With this in mind, we propose to leverage structure prediction models to predict dynamic contact maps of proteins. Structure prediction models introduce a strong inductive bias to learn protein dynamics. This arises from the fact that structural knowledge is required to propose physically coherent hypotheses about the dynamics of the system. Moreover, the evolutionary signals that encode structural information are also likely to encode the dynamics.^10^

While dynamic contact maps represent a coarse view of protein dynamics, many collective variables (CVs) used to represent conformational landscapes in proteins are purely functions of inter-residue distances, since these pairwise distances define the geometry of the backbone. Understanding the dynamic contacts within proteins can yield valuable insights for both experimental and computational studies. For instance, when designing Forster Resonance Energy Transfer (FRET) or Double Electron-Electron Resonance (DEER) experiments, one must carefully select labeling sites that undergo dynamic changes to effectively capture the conformational ensemble of the system.^31,32^ Similarly, mutational studies can benefit from targeting residues involved in dynamic contacts, as these often represent key functional or regulatory regions.^33,34^ In computational research, such as MD simulations, the sheer number of particles makes it challenging to monitor all degrees of freedom. Dynamic contact maps can help define meaningful CVs for enhanced sampling protocols, where an external bias is applied along the CVs,^35^ or for efficient *post hoc* featurization and dimensionality reduction of trajectory data.^36^

In this work, we posit that dynamics can be learned by finetuning structure prediction models, and introduce ESMDynamic, a sequence to dynamic contact map model. We employ ESMFold^8^ as a base model, which is based on ESM2, a PLM that extracts coevolutionary signals without requiring a MSA, and can therefore infer structures without employing expensive sequence alignments, unlike other established methods.^1,3–5^ A two-stage training protocol was applied to fit ESMDynamic, where we first pretrained on experimentally resolved structures and AlphaFold DB^37^ models from 88,294 sequence-aligned clusters (*>* 95% intracluster identity), where structural variations were used to infer dynamic contacts. These clusters comprise 946,397 protein chains from 598,144 uniquely resolved structures, with a median of four conformers per protein. The model was then finetuned on mdCATH,^38^ a dataset of MD simulations covering 5,398 soluble domains, with an average trajectory length of 464 ns. The training objective comprises three tasks: (i) classification of residue–residue pairs as dynamic contacts, distinguishing fluctuating from static or non-interacting pairs; (ii) regression of contact frequency, representing the fraction of time a contact is formed; and (iii) prediction of contact kinetics, assigning coarse-grained timescales for contact formation and dissociation.

Overall, ESMDynamic achieves excellent performance on the sequence-identity aware test split (*<* 20% identity between clusters in train, validation, and test sets), with a balanced accuracy of 80% and recall of 77% when using MD simulations as ground truth. Notably, more than 90% of all positive predictions occur within five residues of true positives, highlighting a strong correlation with the underlying dynamics. Importantly, ESMDynamic surpasses state-of-the-art ensemble prediction models^27,39^ in the classification of dynamic contacts, both in terms of balanced accuracy and inference wall-clock time. This makes ESMDynamic a valuable tool for rapidly predicting dynamic properties in seconds, including features that may be utilized as CVs, obtaining insights that would take weeks or months with MD simulations.

## Results

### Dataset featurization

In this section, we describe the processing of datasets derived from experimental structure clusters (PDB and AlphaFoldDB) and MD simulations (mdCATH). The goal of this featurization step is to construct training targets and input representations for the three prediction tasks addressed by ESMDynamic: dynamic contact classification, contact equilibrium probability regression, and contact kinetics prediction.

Because these targets are derived from heterogeneous data sources with distinct sampling properties, their construction involves several assumptions and design choices. We therefore also discuss the key nuances, comparisons, and limitations associated with dataset featurization.

#### Dynamic contact classification

In this work, we define dynamic contacts as residue pairs that transiently come into spatial proximity using a customary threshold of 8 Å^40^ (see Methods). While dynamic contacts frequently overlap with native contacts, the former are not necessarily a subset of the latter; residue pairs which form short-lived or unstable interactions are also considered dynamic contacts. More specifically, in the mdCATH dataset (320 K), we find that 64.6% of native contacts are also dynamic, while 27.3% of dynamic contacts are also native. For higher temperatures, we observe that the fraction of dynamic contacts that are also native decreases, reflecting the increasing level of structural disorder and the emergence of more non-native interactions (see Table S1).

Moreover, the definition of dynamic contacts involves nuances which are often ignored when defining native contacts. While native contacts are generally defined based on a single static structure, dynamic contacts reflect the intrinsic flexibility of proteins and require considering an ensemble of conformations. In principle, a rigorous definition of dynamic contacts would depend on thermodynamic parameters, such as temperature, that determine the permissible states within the ensemble. ^41^ However, in practice, defining such ensembles is challenging due to variability in the experimental conditions under which protein structures are determined. To address this, we adopt a data-driven approach: for proteins with multiple experimentally determined structures, we identify dynamic contacts as residue pairs whose spatial separation varies above and below the threshold across conformations. A similar approach is applied to MD simulation datasets, where contact variability over time serves as the basis for labeling (see Methods). This framework allows us to systematically annotate dynamic contacts across diverse datasets.

We note that dynamic contacts derived from PDB/AFDB structural clusters (minimum 95% intracluster identity) may involve positions with amino acid substitutions. When aggregating over all residue pairs in the dataset, 5.3% of dynamic contact instances involve at least one mutated residue. When computed per protein and then averaged across proteins, this fraction is 3.1%, indicating that sequence variation contributes to a small subset of dynamic contacts. Mutations present in experimental structures can influence not only contacts directly involving the mutated residue but also distal regions of the protein, particularly in systems exhibiting allosteric coupling. Furthermore, variability in experimental conditions and protocols makes it difficult to ensure that deposited structures correspond to samples drawn from the same thermodynamic ensemble.

Despite these limitations, leveraging heterogeneous structural datasets has proven effective in large-scale protein modeling efforts, including both structure prediction^1–5,8,9^ and studies of conformational variability.^39^ Even when the underlying ensembles are not strictly equivalent, the large number of available structures enables models to learn generalizable structural patterns while averaging over experiment-specific variation, although some biases may remain. In our framework, this issue is mitigated by using structural clusters only during the pretraining stage to expose the model to diverse conformations across many proteins, while the final finetuning stage is performed exclusively on MD simulation data (see Methods), which samples a well-defined thermodynamic ensemble for each protein system. We perform additional characterization of dynamic contacts in Supplementary Results, Section 2.1 (see also Tables S1–3 and Figures S1–5).

#### Equilibrium probability regression

While dynamic contact classification captures whether a residue pair transitions between contact and non-contact states, it does not quantify how frequently these interactions occur within the sampled ensemble. To provide a more complete description of contact behavior, we introduce an additional featurization that estimates the equilibrium contact probability. In this work, we refer to this as the contact frequency (based on a frequentist interpretation of probability), the contact occupancy, or the equilibrium contact probability (we note that thermodynamic equilibrium may not have been reached for all contacts in the training data).

For each residue pair, we compute the fraction of simulation frames in which the contact is formed, which we refer to as the contact occupancy. In a frequentist sense, this quantity approximates the marginal probability of contact formation under the thermodynamic conditions of the simulation. Unlike binary dynamic contact labels, which capture variability, this continuous measure reflects the relative stability of interactions within the sampled ensemble (see Methods).

As with the classification task, these quantities are computed independently for each temperature condition in the mdCATH dataset (320–450 K), enabling the model to learn temperature-dependent changes in contact stability. We emphasize that contact occupancy should be interpreted as an empirical estimate of equilibrium behavior within the accessible sampling window of the MD simulations. The finite trajectory lengths (∼500 ns) may limit convergence for slow conformational processes, and therefore the estimated probabilities primarily reflect the dominant states sampled on nanosecond to sub-microsecond timescales. Despite this limitation, we find that contact occupancy is relatively stable with respect to additional sampling (Table S2), indicating that it provides a robust and informative target for learning.

Taken together with dynamic contact classification, this formulation enables a clear distinction between contact variability (whether a contact forms) and equilibrium contact probability (how often it forms). This separation improves interpretability and avoids conflating intermittency with stability when characterizing residue–residue interactions.

#### Kinetics prediction

While the binary definition of dynamic contacts captures contact variability across sampled conformations, it does not explicitly encode kinetic information. To address this limitation and provide a more complete description of contact behavior, we extend the featurization pipeline to include coarse-grained kinetic targets derived from MD simulations.

Specifically, for each residue pair, we compute two quantities from the mdCATH trajectories: (i) the average time a contact remains in the “on” state before breaking (contact lifetime or on-time), and (ii) the average time required for contact formation (off-time). These quantities are estimated by tracking transitions in the contact time series and aggregating statistics across trajectories (see Methods). Rather than directly regressing continuous timescales, which are inherently noisy and limited by finite trajectory length, we discretize these values into coarse-grained bins spanning the nanosecond regime: always on/off, 1–10 ns, 10–100 ns, 100–300 ns, *>*300 ns, and undefined (transition is not observed). This discretization reflects the finite sampling of the mdCATH dataset, which consists of trajectories of approximately ∼500 ns. Conformational changes occurring on longer (e.g., microsecond to millisecond) timescales are not represented in the training data. We therefore emphasize that the predicted kinetics should be interpreted as coarse-grained estimates within the timescale window accessible to MD simulations, rather than as a complete characterization of protein dynamics.

To characterize the underlying distribution of contact persistence times, we analyzed the empirical distributions of on-times and off-times across the mdCATH dataset (Figure S1).

These distributions are skewed toward relatively short-lived contacts, with most formation and breaking events occurring on the few-nanosecond timescale. This indicates that the dataset is enriched for fast, local fluctuations, with fewer examples of long-lived contacts associated with large-scale conformational transitions.

To further investigate the relationship between contact kinetics and residue-level dynamics, we analyzed the mdCATH dataset using Dyna-1,^42^ a sequence-based predictor of residue flexibility trained on NMR-derived measurements of protein dynamics. Dyna-1 assigns per-residue scores that correlate with the timescale of local motions, providing a complementary perspective on conformational variability. We compared the average contact lifetime for residue pairs as a function of the Dyna-1 scores of the participating residues (Figure S2). We find that contacts involving at least one residue with a high Dyna-1 score tend to exhibit longer lifetimes within the accessible MD timescales (∼1–500 ns). Interestingly, residue pairs in which both residues have high Dyna-1 scores do not show a further increase in lifetime and, in some cases, exhibit slightly reduced lifetimes. We note that these trends are limited to the nanosecond regime captured by the simulations and do not directly probe slower (*µ*s–ms) dynamical processes.

Taken together, these features define a kinetics prediction task that complements the classification and equilibrium probability heads. While the classification head captures contact variability and the frequency head estimates equilibrium occupancy, the kinetics head provides additional information on the timescales of contact formation and persistence.

### Model architecture

ESMDynamic (Figure 1) is built upon the ESMFold framework. ^8^ Structure prediction models offer a powerful prior for inferring protein dynamics, as knowledge of 3D geometry constrains the plausible ways a protein conformation can fluctuate. By leveraging ESMFold, we incorporate structural reasoning directly into the prediction process. An additional advantage of ESMFold is that it extracts evolutionary and structural features from a single input sequence, removing the need for computationally expensive MSAs and enabling high-throughput predictions.

**Figure 1:**
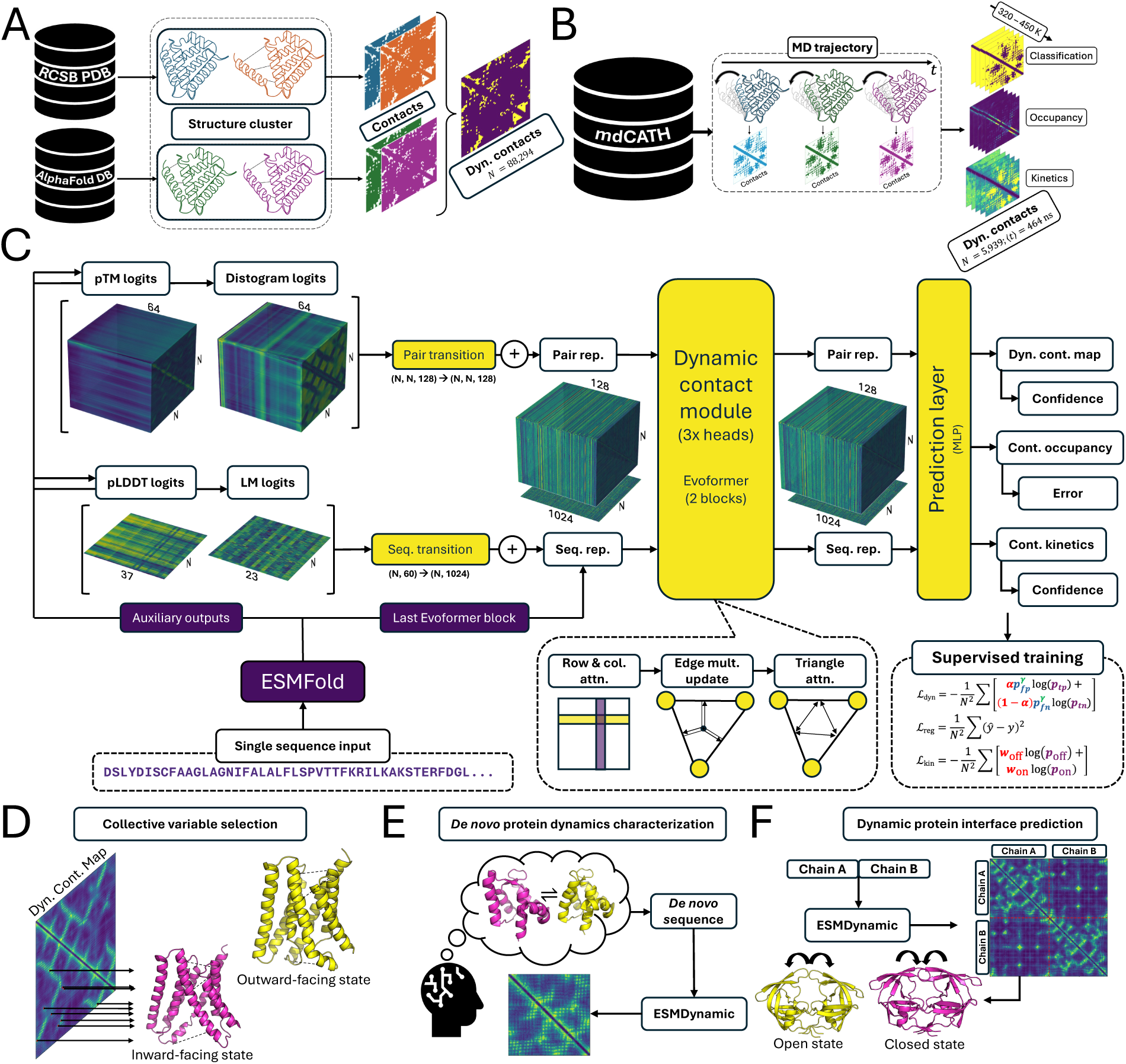
Schematic of the ESMDynamic approach. (A) Sequence-aligned structural ensembles are used to derive dynamic contact maps for pretraining (classification task). (B) Dynamic contacts extracted from MD trajectories (mdCATH) are used for supervised training. (C) Model architecture: a single input sequence is processed by ESMFold; auxiliary outputs are transformed and used to bias the final Evoformer representations, which are passed to Dynamic Contact Modules (DCM) and multilayer perceptrons to predict contact features. The model is trained with focal loss for classification, regression loss for contact frequency, and weighted multiclass cross-entropy for kinetics. (D) Predicted dynamic contacts can be used to define collective variables for MD trajectory analysis or biasing. (E–F) Applications include derivation of collective variables for downstream tasks, prediction of dynamics in *de novo* proteins, and identification of dynamic protein interfaces.

Beyond its core structure prediction capability, ESMFold provides auxiliary signals that are particularly informative for modeling conformational variability. These include distogram logits, which represent predicted distributions of residue–residue distances and can highlight pairs that fluctuate between contact and non-contact states.^1^ Language model (LM) logits provide token-level probabilities that correlate with sequence conservation, often enriched at functionally or structurally important sites.^8^ Uncertainty-related outputs, such as predicted TM-score (pTM) and per-residue confidence (pLDDT), further help distinguish between rigid and flexible regions of a protein.^1^

ESMDynamic integrates these auxiliary signals by biasing the final sequence and pairwise representations from ESMFold’s last Evoformer^1^ block. These enriched features are then processed by three independent *Dynamic Contact Modules* (DCMs), each corresponding to a distinct prediction head with complementary objectives. The Evoformer architecture is well-suited for these tasks, as it models complex, context-dependent relationships between residues using attention-based mechanisms. Table S4 provides parameter counts for different submodules.

The first head performs dynamic contact classification, predicting the probability that a residue pair forms a contact under thermal fluctuations. An auxiliary confidence module estimates the expected accuracy at each sequence position. The second head predicts the contact frequency, defined as the fraction of frames in which a residue pair is in contact, providing an estimate of the equilibrium contact probability. This head is also equipped with an error prediction module that estimates the expected regression error over each pair. Finally, the kinetics head predicts coarse-grained timescale information by classifying both contact formation (off-time) and contact persistence (on-time) into discrete bins spanning the nanosecond regime (see Methods). This head is also equipped with a confidence prediction module that estimates the average accuracy per position. Predictions for all heads are available on the five temperature settings used in mdCATH (320–450 K). Together, these outputs capture contact formation probability, persistence, and coarse-grained kinetics within the timescales accessible to the training data. For details on training see Methods.

### Performance and ablation studies

ESMDynamic was first pretrained on experimentally determined protein structures (classification head) and subsequently finetuned using MD simulations (see Methods). Our validation focuses primarily on the MD dataset. MD trajectories offer a continuous and detailed view of protein conformational variability, whereas experimental structures typically provide only sparse snapshots, limiting their utility for capturing dynamic behavior. This is reflected in the proportion of dynamic contacts resolved by each method; in experimental clusters, dynamic contacts constitute fewer than 1% of all residue pairs, while in MD simulations, this proportion increases to approximately 15% (mdCATH, 320 K), reflecting their greater sensitivity to conformational dynamics (see Supplementary Results, Section 2.1.1).

On the MD-based, sequence-identity-aware test split, ESMDynamic achieves a balanced accuracy of 80% (Figure 2A), with a recall of 77% and a precision of 51% at 320 K (Table S5). Performance for the dynamic contact classification task decreases at higher temperatures (Table S5), consistent with the increased prevalence of transient and disordered interactions. In contrast, the frequency regression head predicts contact occupancy with an RMSE of 7.6% at 320 K, and its performance improves with increasing temperature (Table S6). This trend likely reflects the fact that higher-temperature ensembles exhibit more uniform contact occupancies due to increased disorder, which are easier to approximate. The kinetics classification task is more challenging, resulting in lower overall performance. At 320 K, the model achieves macro recall (equivalent to macro-averaged balanced accuracy) of 34% and 43%, and macro precision of 34% and 47%, for on-time and off-time predictions, respectively (Tables S7–8). Micro-averaged metrics are not reported due to class imbalance, which would otherwise inflate performance estimates. Notably, for a six-class problem, a naive classifier would yield a macro recall of 16.6%, indicating that the model captures meaningful kinetic signal. At higher temperatures, performance shifts toward higher recall at the expense of precision.

**Figure 2:**
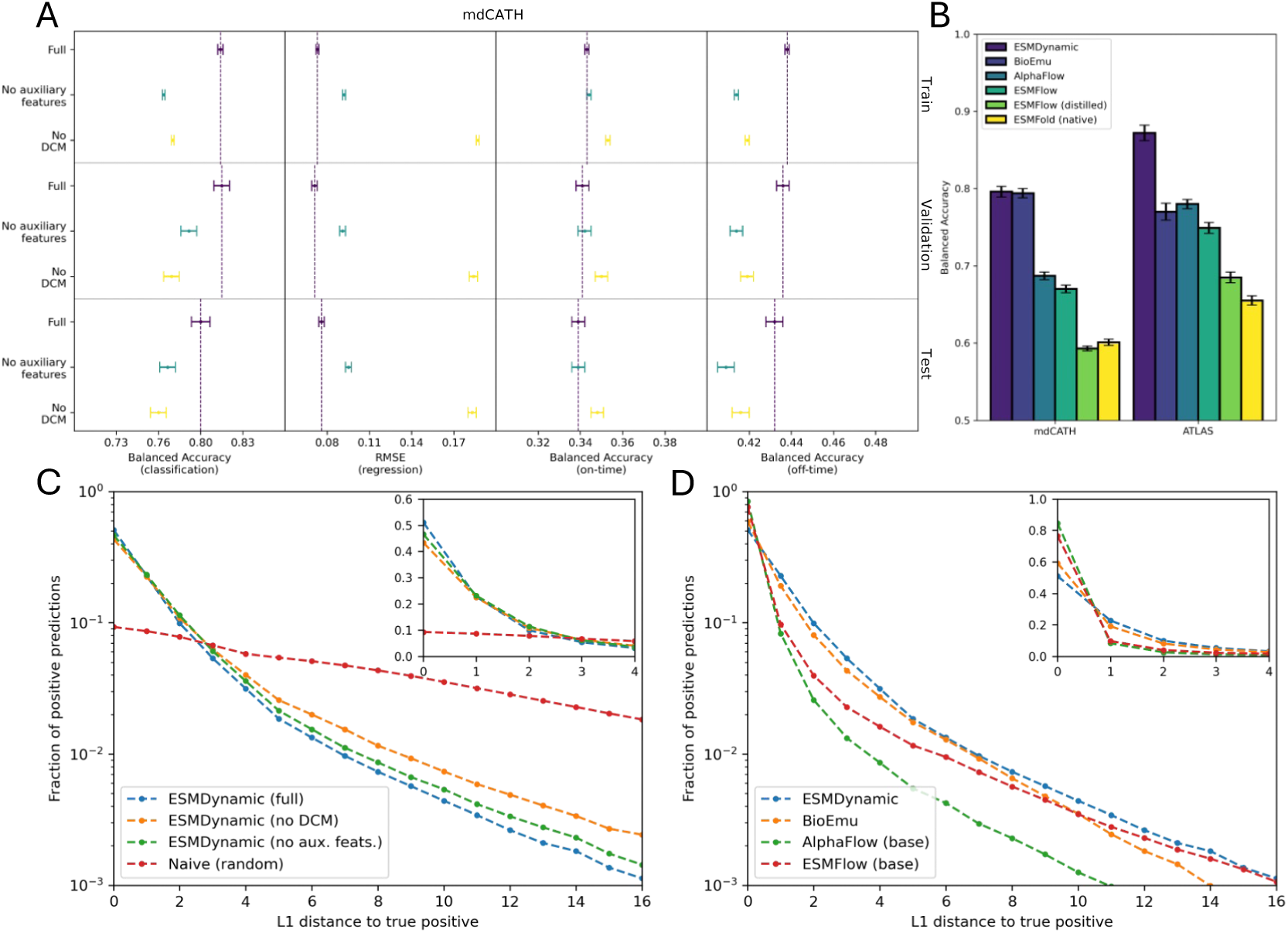
Performance of ESMDynamic, ablated variants, and comparison with baseline and ensemble methods. (A) Balanced accuracy (classification) and RMSE (regression) of the full model and ablated variants across mdCATH (320 K) splits. Error bars denote standard errors. (B) Comparative performance of ESMDynamic, BioEmu, AlphaFlow, and ESMFlow on the mdCATH and ATLAS test sets. Error bars denote standard errors. The naive baseline (ESMFold) assumes all native contacts are dynamic. (C) Fraction of positive predictions as a function of sequence-space (*L*_1_) distance to ground-truth positives for ablated models. A faster decay indicates stronger spatial localization of predictions around true positives, whereas a random baseline yields a more uniform distribution. (D) Same analysis as in (C), comparing ESMDynamic and ensemble methods. High-precision, low-recall models (e.g., AlphaFlow) exhibit steeper decay, reflecting more conservative but spatially concentrated predictions.

We also evaluate the performance of the confidence and error heads on the test set. The dynamic contact confidence head predicts the average per-site accuracy with an RMSE of 8.7% at 320 K. This error increases with temperature, indicating that the model’s confidence estimates become less calibrated at higher temperatures (Table S9). This degradation should be taken into account when interpreting confidence values in high-temperature regimes. For the contact frequency head, the residual RMSE is 5.6% and remains relatively stable across temperatures, suggesting that the residual predictions provide a consistent estimate of uncertainty across conditions (Table S10). In contrast, the kinetics confidence head achieves an RMSE of 10.2% at 320 K, with errors increasing at higher temperatures, again indicating reduced calibration in more disordered regimes (Table S11). In addition, we evaluated the relationship between ESMDynamic uncertainty estimates and ESMFold confidence (pLDDT). Interestingly, we find a strong correlation between these quantities, with Spearman’s *ρ* in the range of 0.7–0.8, where the sign depends on whether the head predicts confidence (positive correlation) or error (negative correlation). These results are shown in Figure S6.

To assess the contribution of different model components, we also evaluated ablated variants lacking auxiliary inputs or the DCM (see Methods). As expected, the removal of the model elements affects its final performance across the training, validation, and test sets (Figure 2A, Tables S12–19). The reduced models generally show marginal to moderate degradation in performance. The ablation results highlight the fact that the features derived from ESMFold can be readily used to predict dynamic contacts, i.e., the model without specialized Evoformer blocks can still attain good performance. However, the prediction task is complex and nuanced: the addition of auxiliary features and a dedicated DCM does improve the performance, indicating the need for additional parameters to correctly predict difficult examples. An interesting exception is the on-time kinetics prediction, where the model with no DCM performs marginally better than the full model, although the difference is less than 1%.

Beyond conventional classification metrics, we also evaluate the spatial correlation between predicted and ground-truth dynamic contacts in sequence space. Specifically, we compute the probability distribution of the shortest L_1_ distance between predicted and true positives (Figure 2C). This distribution’s y-axis intercept is related to precision, and its overall shape captures how tightly the model’s predictions cluster around true dynamic contacts. Figure S7 shows an illustration of how the L_1_ distance is calculated. We find that *>* 90% of positive predictions fall within five residues of a true positive. The full ESMDynamic model shows faster decay than ablated versions, indicating that its dynamic contact predictions are more tightly concentrated around the ground truth compared to the reduced models.

Since we are computing the shortest L_1_ distance to characterize the resolution of the model, we note that even a naive random classifier would show strong results if ground-truth samples were distributed roughly uniformly throughout the map. However, this is not the case, as shown by the probability distribution for the naive random classifier (Figure 2C). To further assess the co-localization of dynamic contacts, we compute their L_1_ radial distribution function (Figure S8), which confirms that dynamic contacts are spatially correlated in sequence space and that the concentration of positive predictions around the ground truth is a feature of such correlation.

### ESMDynamic surpasses ensemble prediction models in accuracy and runtime

We compared ESMDynamic to three state-of-the-art ensemble prediction models: BioEmu,^39^ AlphaFlow, and ESMFlow.^27^ While BioEmu and AlphaFlow are both based on AlphaFold2 and rely on MSAs, ESMFlow is built on the ESMFold architecture. Unlike ESMDynamic, which is specifically trained to predict dynamic contacts, these models are designed to generate full structural ensembles from sequence inputs. They learn to reconstruct conformational variability using both experimental structures and MD simulations. In particular, BioEmu was trained on approximately 200 ms of MD data,^39^ whereas the flow models and ESMDynamic were trained on only a few milliseconds of protein trajectories. Although ensemble-generation models are not explicitly trained to classify dynamic contacts or predict their equilibrium probability, such features can be inferred from their predictions, as dynamic contacts can be directly inferred from structural variability. The models compared here serve as demanding benchmarks, as they have demonstrated superior performance over other ensemble-generation methods, including those based on MSA subsampling.^27,39^ Nonetheless, we note that these models do not generate time-correlated samples, and for this reason we cannot compare them against ESMDynamic in the kinetics prediction task.

When evaluated on dynamic contact prediction, ESMDynamic matches or substantially outperforms all benchmarks (Figure 2B). ESMDynamic and BioEmu achieve similar balanced accuracy on the mdCATH test set. Notably, a large portion (46 ms) of BioEmu’s training data was derived from CATH sequences,^39^ so overlap between the test set used in this work and BioEmu’s training data cannot be ruled out. In contrast, the flow-based models tend toward high-precision, low-recall predictions, resulting in lower overall balanced accuracy. Specifically, on the mdCATH test set, AlphaFlow achieves the highest precision (84%), but with low recall (40%), yielding a balanced accuracy of 69% averaged across proteins. While BioEmu does not attain the top balanced accuracy, recall, or precision, it achieves the highest F1 score (60%), compared to 57% for ESMDynamic, as well as the lowest contact occupancy RMSE (6.3% vs. 7.6% for ESMDynamic). See Table S20 for a full comparison.

All models were also evaluated on the AlphaFlow test split derived from ATLAS.^43^ Although mdCATH and ATLAS are both MD datasets, they differ in key parameters, including force field choice and simulation temperature^38,43^ (see also Supplementary Results, Section 2.1.2). Notably, the flow models were trained on ATLAS ensembles, whereas ESMDynamic had not been exposed to this data during training. Despite this, ESMDynamic significantly outperforms the flow models and BioEmu in identifying dynamic contacts, achieving a balanced accuracy of 87% (Figure 2B). While BioEmu was not trained on ATLAS data, it performs comparably to the best-performing flow model, AlphaFlow. Additional performance metrics for all models are provided in Table S21. Nonetheless, the generative models exhibit higher precision (e.g., AlphaFlow at 68%) and lower contact occupancy RMSE (e.g., AlphaFlow at 4.4%). Finally, analysis of the minimum L_1_ distance distribution (see previous section) shows faster decay for these models compared to ESMDynamic, consistent with their preference for precision over recall (Figure 2D).

The relatively lower balanced accuracy of the flow models is likely a consequence of their design, in which sampling is constrained by the AlphaFold2 or ESMFold structure module (i.e., noised inputs are consistently passed through the structure module to generate final conformations, rather than using independently trained generative weights). These constraints favor the generation of physically plausible, low-energy conformations, but limit the ability to represent low-probability states in which contacts are broken. In contrast, ESMDynamic is trained directly to predict dynamic contact probabilities without enforcing strict geometric consistency. As a result, it can assign high probabilities to multiple, potentially incompatible contact states, better reflecting the range of structural fluctuations.

We also evaluated a naive baseline that labels native contacts predicted by ESMFold as dynamic. While this baseline performs comparably to the distilled version of ESMFlow on mdCATH, it fails to capture the complexity of the dynamic contact landscape, leading to poor balanced accuracy. To estimate the upper bound of this approach, we additionally used ground-truth native contacts derived from PDB structures. Under this idealized setting, we find that ESMFold performs close to the maximal achievable balanced accuracy (61% vs. 63% for mdCATH at 320 K; see Table S22).

For completeness, we also compared ESMDynamic against BioEmu on four curated collections of experimental structures originally used as validation sets in BioEmu.^39^ In this work, these datasets are repurposed for the dynamic contact classification task and are therefore not evaluated using the original metrics. We note that some structures in these sets may overlap with the ESMDynamic pretraining data. The results of this analysis are presented in Supplementary Results (Section 2.2) and Table S23.

To further interpret and calibrate the nature of the dynamic contacts predicted by ESMDynamic, we ran our model on RelaxDB,^42^ a 133-protein dataset containing residue labels corresponding to characteristic timescales of motion based on NMR data. The analysis of these predictions is presented in Supplementary Results (Section 2.3) and Figure S9.

In terms of runtime, ESMDynamic predicts dynamic contact maps in just a few milliseconds to a few minutes, depending on protein length, since it requires only a single forward pass. Its runtime is virtually identical to that of ESMFold. In contrast, generative methods require up to hundreds of forward passes per protein to sample enough structures, significantly increasing computation time. Among the generative models compared, the distilled version of ESMFlow was the fastest, albeit at the cost of lower accuracy. A timing analysis is presented in Figure S10. AlphaFlow was excluded from this benchmark due to its substantially higher computational cost relative to the other methods, rendering it non-competitive in terms of speed. Overall, ESMDynamic is consistently faster across the full range of evaluated protein lengths.

In addition to evaluating our model against ablated variants and generative models, we also examine the correlation between ESMDynamic’s classification head predictions and the unsupervised contact probabilities produced by ESM2.^8^ Similar to the comparison against the naive baseline in Figure 2B, this test serves to assess whether ESMDynamic is simply recovering native contact patterns rather than learning features specific to conformational dynamics. If the model’s output were highly correlated with ESM2’s unsupervised contact predictions, it would suggest that the dynamic contact probabilities reflect static structural features rather than dynamic behavior. However, on the mdCATH test set (320 K), we observe no meaningful correlation between ESMDynamic’s predictions and those from ESM2 (Figure S11). This supports the conclusion that ESMDynamic is learning non-trivial, dynamics-specific signals that are distinct from those captured by unsupervised models trained solely on sequence data.

### ESMDynamic recovers experimental dynamics in proteins beyond its training set

Figure 3 illustrates the application of ESMDynamic to the neutral amino acid transporter ASCT2/SLC1A5, which is upregulated in cancer and also serves as a retroviral receptor.^44^ The ASCT2 sequence shares less than 20% identity with proteins in both the pretraining and finetuning datasets (see Methods). Although ESMDynamic was pretrained on diverse structural ensembles, it was subsequently finetuned on the mdCATH database, which lacks membrane proteins. This raises the question of whether the model can retain accuracy on transmembrane transporters, making ASCT2 a stringent out-of-distribution test case.

**Figure 3:**
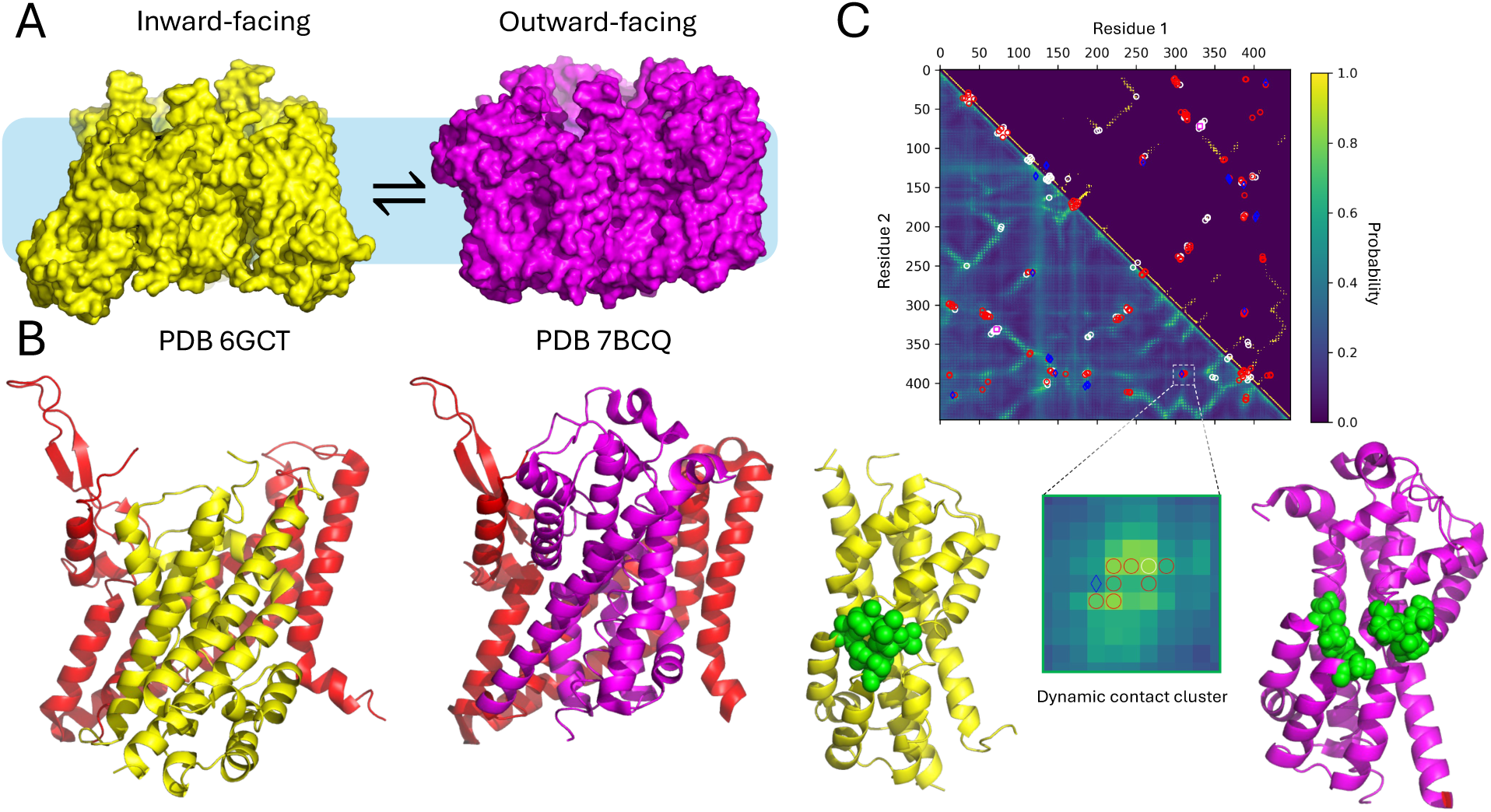
Sample output from ESMDynamic for the ASCT2/SLC1A5 transporter. (A) Experimental structures of the inward-facing and outward-facing conformational states. (B) A cartoon representation highlights a structurally invariant region (red) and regions that undergo conformational change (yellow and purple, corresponding to each state). (C) Dynamic contact map predicted by ESMDynamic (lower triangle) and native contacts predicted by ESMFold (upper triangle). Residue indexing begins at D43. Annotations denote 149 dynamic contacts inferred from the experimental structures. White circles: dynamic contacts predicted by both ESMDynamic and native contacts baseline (62/149). Red circles: dynamic contacts predicted by ESMDynamic but not native contacts baseline (73/149). Magenta squares: dynamic contacts predicted by native contacts baseline but not ESMDynamic (1/149). Blue diamonds: dynamic contacts missed by both ESMDynamic and native contacts baseline (13/149). The inset shows a zoomed-in view of a dynamic contact cluster involving residues S351 to A355 and G430 to P432. These contacts are present in the inward-facing state but are disrupted during the transition to the outward-facing state. Residues involved in these interactions are highlighted in green.

To evaluate performance, we analyzed two experimental structures corresponding to inward-facing (PDB 6GCT)^45^ and outward-facing (PDB 7BCQ)^46^ conformations, which define a set of experimentally validated dynamic contacts. In Figure 3C, we directly compare ESMFold native contacts (upper triangle) and ESMDynamic predictions (lower triangle), while marking ground-truth dynamic contacts with symbols indicating which model correctly identifies each interaction.

Out of 149 experimentally inferred dynamic contacts, ESMDynamic correctly detects 135 (recall of 90%), whereas the native-contact baseline derived from ESMFold detects only 63 (recall of 42%). Overall, ESMDynamic achieves a balanced accuracy of 93%, substantially outperforming the ESMFold baseline (70%). Precision is low for both models (3.0% for ESMDynamic and 4.6% for ESMFold), reflecting the extreme sparsity of dynamic contacts derived from a limited number of experimental structures (see Supplementary Results, Section 2.1.1). Despite this, the comparison clearly shows that the majority of experimentally observed dynamic contacts are uniquely captured by ESMDynamic and are absent from the native-contact baseline.

Interestingly, ESMDynamic (but not the baseline) identifies a discrete dynamic contact cluster (Figure 3C, inset) between residues S351–A355 of hairpin 1 (HP1) and G430–P432 of hairpin 2 (HP2). This interface is functionally important, as HP2 constitutes both the extra-cellular and intracellular gate^47^ and must move to admit or expel the substrate. Consistent with this mechanism, the HP1–HP2 interface is disrupted in the *apo* outward-facing state (PDB 6MP6^48^) and upon binding of the proline-derived inhibitor Lc-BPE at S353 (PDB 7BCQ),^46^ which stabilizes the outward-facing conformation. In contrast, this interface is preserved in the inward-facing conformation (PDB 6GCT).^45^ These observations indicate that ESMDynamic captures functionally relevant, state-dependent interactions that are not encoded in static native contacts.

In summary, ESMDynamic successfully detects key dynamic features governing the transition between inward- and outward-facing states in a transmembrane transporter with minimal sequence similarity to the training set. The explicit comparison against the native-contact baseline highlights that dynamic contacts cannot be recovered from static structures alone, as most arise from transient, non-native interactions. The ability of ESMDynamic to generalize to membrane proteins, despite their absence in mdCATH, suggests that finetuning does not degrade performance in this class of systems. While ESMDynamic itself was not trained on this transporter, ESMFold likely provides a strong structural prior through homologous sequences, enabling the model to build upon accurate structural features while learning dynamics-specific signals. Overall, this case study demonstrates that ESMDynamic can provide meaningful insights into clinically relevant proteins and generalizes beyond its training distribution.

### Dynamic contact maps enable automatic collective variable selection for MD simulations

A key application of ESMDynamic is leveraging predicted dynamic contact maps to identify relevant CVs for analysis or biasing in MD simulations. In this setting, users examine predicted dynamic contacts and employ either automated tools or knowledge-based criteria to select informative metrics to monitor. While dynamic contact maps highlight residue pairs expected to undergo distance changes, their utility extends beyond purely distance-based features. For instance, an interaction may break due to torsional changes within residues, and clusters of dynamic contacts can signal larger conformational rearrangements. In this section, we demonstrate an automated pipeline for CV selection using 145 *µs* of unbiased MD simulations of *Oryza sativa* (rice) SWEET2b,^49^ a transporter that facilitates passive sugar transport along concentration gradients with relevance to crop yield.^50^ Notably, this protein was excluded from the ESMDynamic training set, with no homologs above 20% sequence identity present during training. This ensures the analysis reflects a realistic use case, where ESMDynamic is applied to a previously unseen protein sequence lacking close homologs in the training data.

The proposed pipeline is not intended to prescribe a fixed procedure for utilizing ESMDynamic predictions. Rather, it demonstrates that a simple and fast analysis of predicted dynamic contacts can effectively identify meaningful CVs for MD trajectory analysis. As shown in Figure 4A, the pipeline begins by filtering residue–residue pairs to retain only high-order dynamic contacts. Specifically, predicted positive contacts between residues separated by at least 40 positions in sequence space are selected. This step removes local dynamic contacts (e.g., those within or between adjacent transmembrane helices).

**Figure 4:**
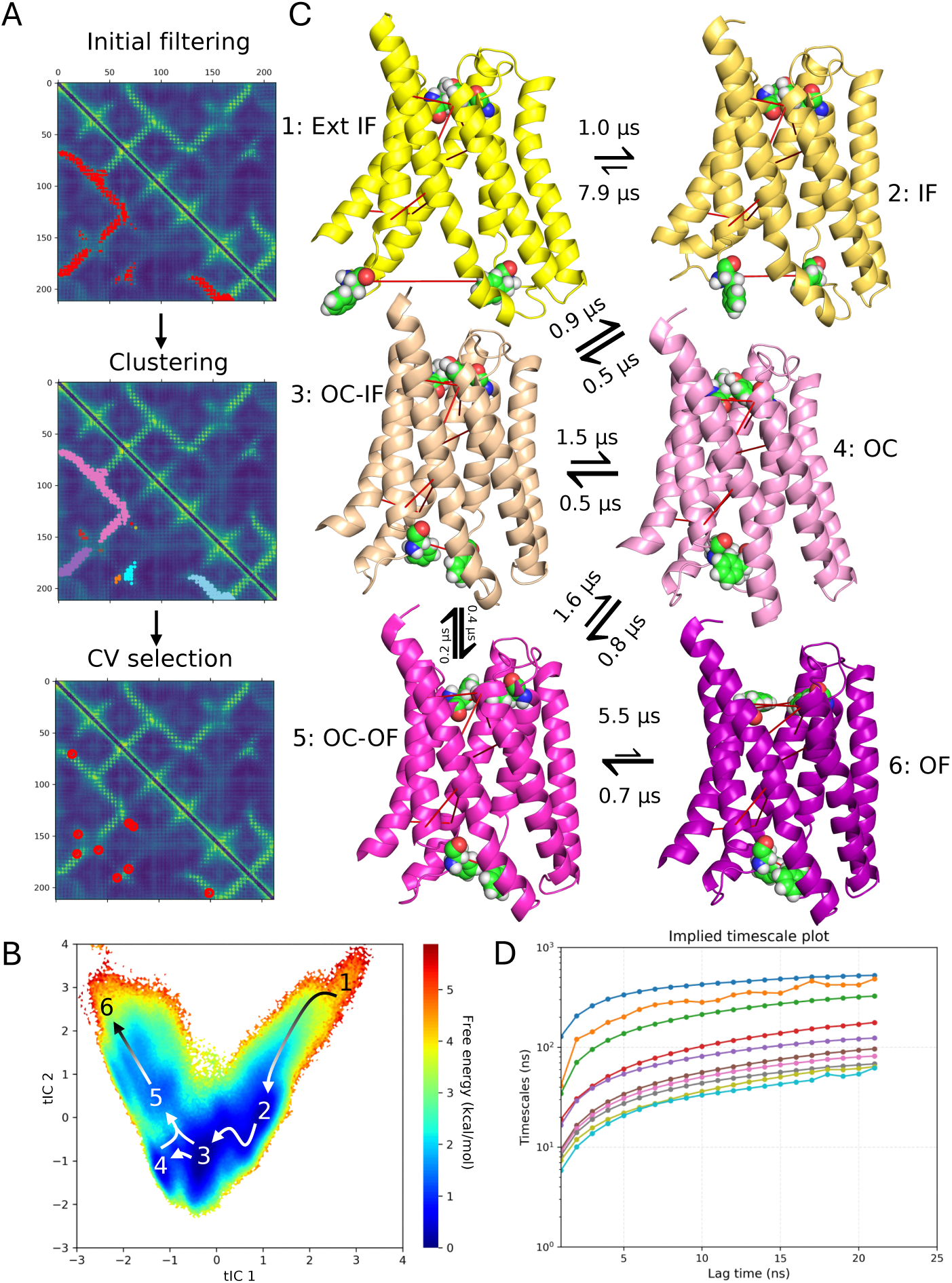
Automatic CV selection with ESMDynamic and resulting MSM analysis for SWEET2b. (A) CV selection pipeline: predicted dynamic contacts are filtered for long-range pairs, clustered into dynamic regions, and representative residue pairs are selected as CVs. (B) Free energy landscape projected onto the top two tICA components, revealing a continuous transition pathway connecting six metastable states. Arrows indicate dominant transitions inferred from the MSM. (C) Representative structures corresponding to the identified states: extended inward-facing (Ext IF), inward-facing (IF), occluded inward-facing (OC-IF), occluded (OC), occluded outward-facing (OC-OF), and outward-facing (OF). Selected CVs are shown as red lines. Molecular gates (F42–F164 and Y60–F191) are shown in space-filling representation. Transition timescales between states are indicated. (D) Implied timescale plot demonstrating convergence of slow dynamical processes.

Next, hierarchical clustering is applied to the remaining residue pairs to identify distinct dynamic regions within the protein. We treat contact map positions as spatial “coordinates” and perform single-linkage clustering using L_1_ distances via Scipy,^51^ limiting the number of clusters to 9 to enable fair comparison with manually curated CVs.^49^ From each cluster, we select the highest-probability dynamic contact and track the corresponding inter-residue distances as CVs.

These distances are then used to featurize the MD trajectory, followed by time-lagged independent component analysis (tICA)^52,53^ for dimensionality reduction. Markov State Models (MSMs) are subsequently constructed from the tICs using the same protocol as in ref. 49 (see also Supplementary Methods). tICA identifies the slowest decorrelating collective motions in the trajectory by maximizing autocorrelation over a specified lag time. This provides a reduced set of kinetically relevant coordinates that are particularly well suited for constructing MSMs. An MSM represents the conformational space as a set of discrete states and estimates the probabilities of transitioning between them over a given lag time. This framework enables the reconstruction of long-timescale dynamics from shorter MD trajectories by exploiting the Markovian property of memoryless transitions.^54^

Figure 4B shows the free energy landscape projected onto the top two tICA components, revealing a continuous transition pathway connecting multiple metastable states. Six representative states are identified (Figure 4C), corresponding to extended inward-facing (Ext IF), inward-facing (IF), occluded inward-facing (OC-IF), occluded (OC), occluded outward-facing (OC-OF), and outward-facing (OF) conformations. The dominant transitions between these states, along with their associated timescales, are captured by the MSM and reflect a sequential alternating-access mechanism. Importantly, the lowest free energy regions correspond to inward-facing and occluded conformations, while the extended-IF and OF states appear as higher free energy, transient states. This is consistent with the *apo* (substrate free) simulation conditions, in which the outward-facing conformation is only transiently stabilized to prevent reverse transport.^49^ Figure 4C also shows the selected CVs projected onto representative structures obtained via MSM clustering, highlighting residue pairs associated with intracellular (F42–F164) and extracellular (Y60–F191) gating motions. These interactions constitute central dynamic features of SWEET2b, consistent with an alternating-access mechanism in which one gate opens to permit substrate binding while the other subsequently opens to enable release. Both thermodynamic and kinetic properties are in strong agreement with those obtained from expert-curated CVs. Specifically, we observe an energy barrier of ∼4 kcal/mol between the IF and OF states, consistent with prior work.^49^ Transition path theory (TPT)^53^ analysis indicates that the IF → OF cycle occurs on a timescale of approximately 6 *µ*s, with the rate-limiting step corresponding to the OC-OF → OF transition, also consistent with previous findings. Additionally, we observe direct transitions between OC-IF and OC-OF states that bypass the fully occluded state, in agreement with prior studies.

To assess the quality of the selected CVs, we compare MSMs constructed from ESMDynamic-derived features against those obtained using expert-selected CVs, as well as baselines derived from MD-based dynamic contacts and ESMFold native contacts using the same pipeline (see Supplementary Methods for details). We use standard MSM validation metrics. The implied timescales (Figure 4D) show convergence at the selected lag time (see Supplementary Methods), and the Chapman–Kolmogorov test confirms Markovian consistency (Figure S12). The MSM constructed from ESMDynamic-selected CVs achieves a VAMP-2 score^55^ of 89% of the theoretical maximum, surpassing the literature value (86%) and outperforming both MD-derived CVs (85%) and ESMFold native contact CVs (84%). All VAMP-2 scores were computed using the six slowest processes to match the methodology of ref. 49.

Beyond this quantitative improvement, ESMDynamic-selected CVs recover the same qualitative transition mechanism as the expert-defined CVs, identifying the same sequence of metastable states and transition pathways across the transport cycle. In contrast, baseline methods yield free energy landscapes that deviate both qualitatively and quantitatively (Figure S13). In particular, CVs derived from ESMFold native contacts fail to capture key functional motions, notably missing important gating interactions, resulting in MSMs that do not resolve the full transport cycle. While CVs derived from MD-based dynamic contacts correctly identify both gating motions, the clustering-based selection introduces variability that leads to distortions in the reconstructed free energy landscape relative to the expert reference. This discrepancy is evident in Figure S13C–E, where alternative CV selections fail to reproduce the transition topology observed with ESMDynamic and expert-defined CVs. Overall, these results demonstrate that ESMDynamic enables identification of mechanistically meaningful CVs directly from sequence, producing MSMs that accurately reproduce both the thermodynamic landscape and kinetic pathways of the system. Notably, this level of agreement with expert-designed CVs is achieved without access to trajectory data, highlighting the ability of ESMDynamic to extract functionally relevant degrees of freedom from sequence alone.

### ESMDynamic captures conformational switching in a *de novo* design

Recent advances in deep learning have enabled the design of multistate proteins that can dynamically transition between distinct conformations.^56–58^ These systems provide a stringent benchmark for evaluating ESMDynamic, as they test the model’s ability to generalize principles of conformational dynamics to sequences that are entirely absent from natural evolutionary space. One such example is a newly engineered variant of the N-terminal domain of troponin C, a calcium-binding protein designed to adopt two alternative conformations under standard thermodynamic conditions. Upon calcium binding, the equilibrium shifts between these states.^58^ Because this protein was deposited after ESMDynamic training and is *de novo* designed, it represents a particularly challenging out-of-distribution test case. Moreover, unlike earlier design efforts that emphasize large-scale interdomain rearrangements or rigid-body motions,^57^ this system undergoes intradomain conformational changes, making it well suited to assess sensitivity to subtle contact rearrangements.

For this case study, we analyzed the I89 variant.^58^ In Figure 5A, we directly compare ESMFold native contacts (upper triangle) and ESMDynamic predictions (lower triangle), while marking experimentally derived dynamic contacts from *apo* structures (PDB IDs: 9CIG and 9CID) with symbols indicating prediction outcomes. ESMDynamic correctly identifies 47 out of 49 dynamic contacts (recall of 96%), whereas the native-contact baseline recovers only 18 (recall of 37%). Overall, ESMDynamic achieves a balanced accuracy of 88%, substantially higher than the ESMFold baseline (66%). Precision remains low for both models (6% for ESMDynamic and 9% for ESMFold), which we attribute to the extreme sparsity of dynamic contacts derived from a limited number of experimental conformations (see Supplementary Results, Section 2.1.1). Importantly, the majority of experimentally observed dynamic contacts are uniquely captured by ESMDynamic and are not present in the native-contact baseline.

**Figure 5:**
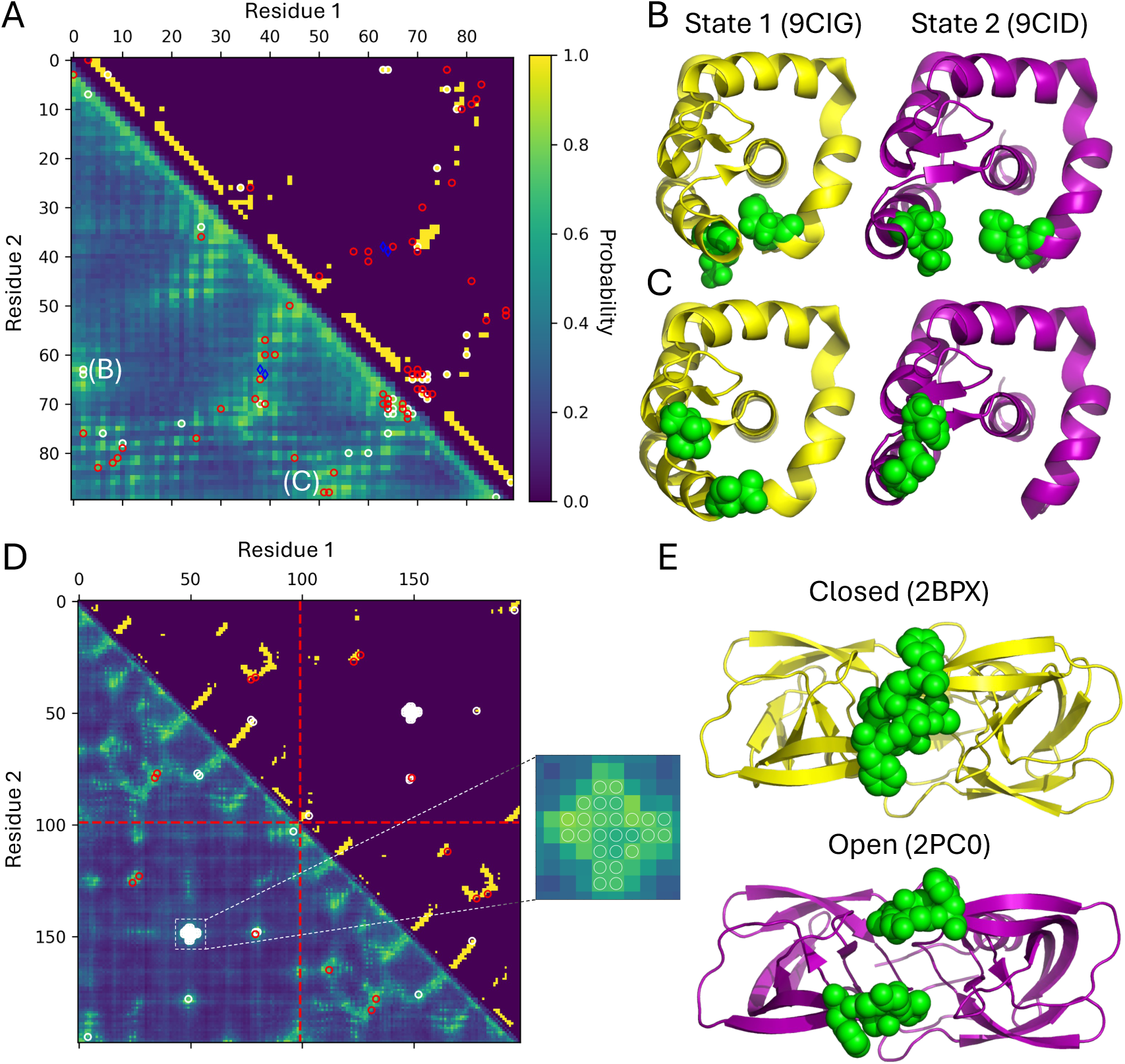
Additional ESMDynamic applications. (A) Predicted dynamic contact map (ESMDynamic, lower triangle) and native contact map (ESMFold, upper triangle) for *de novo* multistate design “I89” from troponin C (residues indexed from A5). White circles: dynamic contacts predicted by both ESMDynamic and native contacts baseline (18/49). Red circles: dynamic contacts predicted by ESMDynamic but not native contacts baseline (29/49). Blue diamonds: dynamic contacts missed by both ESMDynamic and native contacts baseline (2/49). Contacts M7–K68 and K44–G75 are labeled B and C. (B) Experimental states 1 and 2 with M7 and K68 in green. (C) Experimental states 1 and 2 with K44 and G75 in green. (D) Predicted dynamic contact map (ESMDynamic, lower triangle) and native contacts (ESMFold, upper triangle) for HIV-1 protease homodimer. Residues indexed from P1; chains indexed consecutively. White circles: dynamic contacts predicted by both ESMDynamic and native contacts baseline (34/42). Red circles: dynamic contacts predicted by ESMDynamic but not native contacts baseline (8/42). Red dashed line: chain boundary. Inset shows a dynamic contact cluster at the dimer interface (I47–F53), identified by ESMDynamic as a high-probability site (also native contacts). (E) Experimental structures with I47–F53 in green; top: closed conformation, bottom: open.

Beyond recovering known interactions, ESMDynamic predicts additional high-probability contacts in nearby regions. In particular, we observe a probability-enriched segment spanning the distal portion of *α*-helix A (residues A29–F33), a flexible loop (D34–G38), and *α*-helix B (Y43–L53). Although these regions appear structurally similar across the two experimental states, their proximity to flexible loop elements suggests that local fluctuations give rise to transient contact formation and breaking. This result is consistent with Figure S15 of ref. 58.

Figures 5B and 5C further illustrate representative dynamic contacts in this system. The residue pair M7–K68 corresponds to a contact favored in state 1 and is consistent with a native state contact, whereas the pair K44–G75 is favored in state 2 and is not captured by native contacts. ESMDynamic correctly identifies both interactions, demonstrating its ability to capture state-dependent contact switching, including interactions that are absent from the ESMFold static structure.

In summary, ESMDynamic accurately recovers experimentally observed dynamic contacts in a *de novo* multistate protein and identifies additional candidate interactions associated with local flexibility. The explicit comparison with the native-contact baseline highlights that many of these contacts arise from transient, non-native interactions, which cannot be inferred from static structure prediction alone. These results demonstrate that ESMDynamic generalizes beyond natural proteins and can capture subtle intradomain conformational changes in designed systems.

### ESMDynamic shows emergent ability to predict dynamic protein interfaces

Although ESMDynamic was not explicitly trained on multimeric proteins or protein interfaces, it inherits inductive biases from its base model, ESMFold, which has demonstrated emergent capabilities for multimer structure prediction despite lacking multimer-specific supervision.^8^ We refer to this behavior as “emergent” because it arises without direct training on multimeric systems. In this context, an important question is whether ESMDynamic can similarly identify dynamic contacts at protein–protein interfaces.

A systematic evaluation of interface dynamics remains challenging due to limited data availability, particularly the scarcity of multimeric systems resolved in multiple conformational states or sampled extensively via MD simulations. To address this, we present a representative case study on a well-characterized homodimer, the HIV-1 protease. This viral enzyme plays a central role in HIV maturation and is a major target in antiretroviral therapy.^59,60^ Its function depends critically on dimer interface dynamics, particularly in the region spanning residues I47 to F53, which forms part of the binding pocket gate and undergoes conformational changes to accommodate inhibitors.^61^

Leveraging experimentally resolved structures in both closed^62^ and open^63^ conformations, we define a set of interface dynamic contacts and evaluate model predictions. In Figure 5D, we directly compare ESMFold native contacts (upper triangle) and ESMDynamic predictions (lower triangle), with symbols indicating which model correctly identifies each experimentally derived dynamic contact. ESMDynamic recovers all dynamic contacts (recall of 100%), whereas the native-contact baseline achieves 81% recall. ESMDynamic attains a balanced accuracy of 96%, compared to 89% for the baseline. While the baseline performs relatively well in this system due to extensive overlap between native and dynamic contacts, ESMDynamic still provides a consistent improvement and captures all experimentally observed interface rearrangements. As in previous case studies, precision is low for both models (3% for ESMDynamic and 6% for ESMFold), reflecting the sparsity of dynamic contacts derived from a small number of experimental conformations (see Supplementary Results, Section 2.1.1).

Beyond the experimentally validated contacts, ESMDynamic predicts additional high-probability interactions within the dimer interface. Some of these are not observed in the available crystal structures, which we attribute to the limited sampling of only two conformational states. In contrast, ESMDynamic is trained on MD-derived ensembles and can therefore capture transient interactions that may arise under physiological conditions but are not resolved experimentally. The model also identifies intrachain dynamic contacts, further suggesting that it captures both inter- and intra-molecular flexibility within the system.

In summary, ESMDynamic exhibits emergent capabilities analogous to those of ESMFold, extending them from static structure prediction to dynamic contact inference at protein interfaces. While comprehensive benchmarking remains limited by data availability, this case study demonstrates that ESMDynamic can identify functionally relevant interface dynamics in multimeric systems, provided that a reliable structural model of the complex is available via ESMFold.

### ESMDynamic predicts dynamic contacts at the proteome scale

A key advantage of ESMDynamic over generative approaches is its computational efficiency, which enables large-scale application to proteome-wide datasets. To demonstrate this capability, we applied ESMDynamic to the human proteome (UniProt Proteome ID UP000005640). Due to hardware constraints (NVIDIA RTX 4090 GPU with 24 GB VRAM), we imposed a sequence length cutoff of 1000 residues. This threshold still covers approximately 88% of proteins in the human proteome (18,238 out of 20,659 proteins; see Figure S14A for the length distribution). All predictions were performed for monomeric chains. For completeness, we also provide the corresponding ESMFold-predicted structures alongside ESMDynamic outputs in our data repository.

Proteome-scale inference required approximately 48 GPU-hours on a single RTX 4090, demonstrating that ESMDynamic enables practical, large-scale screening of conformational variability on consumer-grade hardware. The computational cost is orders of magnitude lower than that required for generating comparable conformational ensembles using MD simulations or generative models, making ESMDynamic well-suited for high-throughput applications such as identifying flexible regions, prioritizing targets for simulation, or annotating large protein datasets.

In addition to scalability, we observe that the confidence head outputs of ESMDynamic correlate with per-residue confidence scores (pLDDT) from the corresponding ESMFold predictions (Figure S14B–D). This suggests that uncertainty in dynamic-contact predictions is linked to structural uncertainty, providing a consistent and interpretable signal for downstream analysis.

Together, these results establish ESMDynamic as a practical tool for proteome-scale characterization of protein dynamics, enabling rapid identification of conformationally variable regions across large sequence datasets.

## Discussion

ESMDynamic provides a framework for predicting dynamic residue–residue contacts directly from sequence. Unlike approaches that rely on explicit ensemble generation, the model outputs residue-pair probabilities that reflect contact variability under the conditions represented in the training data. This representation offers a compact description of conformational flexibility that can be used in downstream structural and simulation-based analyses.

Beyond binary contact variability, ESMDynamic incorporates multiple prediction heads that capture complementary aspects of contact behavior. In addition to the classification head, which estimates the probability of contact formation, the model includes a contact frequency head that predicts the fraction of frames in which a contact is formed, and a kinetics head that assigns coarse-grained timescale bins for contact formation and breaking. Predictions are provided across the temperature range represented in the mdCATH dataset. Together, these outputs extend the representation from simple contact variability to include contact probability, persistence, and approximate kinetic behavior within the accessible timescales and conditions of the training data.

Our benchmarks show that ESMDynamic performs competitively with state-of-the-art ensemble generation models (BioEmu, AlphaFlow, and ESMFlow) for the task of dynamic contact prediction. We attribute this to the use of direct supervision on dynamic contact labels, which enables the model to learn sequence-dependent features associated with contact variability. In contrast, flow-based models are constrained by their underlying structure modules, which bias sampling toward high-likelihood conformations. This can limit their ability to represent low-probability states in which contacts are disrupted, leading to higher precision but lower recall. ESMDynamic, by design, does not enforce strict structural consistency at the prediction stage, allowing it to capture a broader set of potential contact states. At the same time, BioEmu achieves comparable performance on several metrics, highlighting that different modeling strategies involve distinct trade-offs.

Throughout this work, the primary “naive” baseline for dynamic contact prediction is given by native contacts derived from a single predicted structure. A natural extension is to consider whether uncertainty-related outputs from structure prediction models, such as predicted aligned error (PAE) or distogram distributions, could provide stronger baselines by identifying flexible or ambiguous regions. While this approach is conceptually appealing and has been explored in prior work, available evidence suggests important limitations. In particular, distogram-derived distance distributions often exhibit multimodal or irregular behavior^64^ that does not consistently correspond to physically meaningful conformational ensembles, as highlighted by comparisons to experimental distance measurements (e.g., DEER spectroscopy^65^). As a result, simple heuristics that interpret multimodality or uncertainty as indicators of contact formation and breaking may be unreliable without additional modeling assumptions. In contrast, ESMDynamic is trained directly on MD-derived contact variability, enabling it to learn representations that better reflect the patterns present in these datasets. For this reason, we focus our comparisons on generative ensemble models, which provide a more appropriate baseline for conformational variability.

ESMDynamic shows evidence of generalization beyond its training distribution in the case studies considered here. Although the model was not explicitly trained on designed proteins or multimeric systems, it recovers experimentally supported dynamic contacts in these examples. This behavior is inherited in part from the structural priors encoded in ESMFold. However, these results should be interpreted cautiously, as they are based on a limited number of case studies rather than systematic benchmarks.

The predicted dynamic contact maps can be used to support downstream analyses. In the SWEET2b example, they enable automated selection of collective variables for MSM construction using only sequence information. The resulting MSMs recover key conformational states and achieve performance comparable to those obtained using expert-selected features. This suggests that dynamic contact predictions can provide a useful starting point for identifying relevant degrees of freedom, although they do not replace detailed analysis of simulation data.

In addition to predictive performance, ESMDynamic has practical advantages in terms of computational cost. Because the model requires only a single forward pass, predictions can be generated in seconds to minutes per protein, depending on sequence length. In contrast, ensemble-based generative approaches require sampling large numbers of structures, resulting in substantially higher computational expense. This difference makes ESMDynamic well suited for large-scale or exploratory applications.

Several limitations should be considered when interpreting these results. The training data are derived primarily from MD simulations of ordered, monomeric proteins, which biases the model toward this regime. As a result, performance degrades for intrinsically disordered proteins (see Supplementary Results, Section 2.4 and Figure S15) or systems with low structural confidence, where the underlying ESMFold predictions are uncertain (see Figure S7). In addition, the MD datasets used for training are limited in timescale (typically hundreds of nanoseconds), which constrains the range of dynamical processes that can be learned. Consequently, ESMDynamic is expected to preferentially capture faster, local fluctuations rather than slower, large-scale conformational transitions.

More broadly, the predicted quantities provide a coarse-grained summary of conformational variability and do not directly encode detailed mechanistic information and may conceal global geometric shifts. While they can highlight regions of interest and suggest candidate interactions, they must be combined with other approaches, such as MD simulations, enhanced sampling methods, or experimental measurements, to obtain detailed mechanistic insight. In this context, ESMDynamic is best viewed as a complementary tool that can guide and prioritize more detailed analyses.

In summary, ESMDynamic demonstrates that dynamic contact prediction from sequence is feasible using supervised learning on MD-derived data. The model provides a fast and scalable way to approximate aspects of conformational variability, with potential applications in protein analysis and simulation workflows, while remaining subject to the limitations of the training data and representation.

## Methods

### Datasets

#### RCSB and AlphaFold DB structure ensembles

Structural ensembles were sourced from the RCSB Protein Data Bank (PDB). Both experimental structures and AlphaFold DB^37^ models were utilized. MMseqs2-based groupings^6,7^ were used to generate 128,660 clusters with *>*95% intra-cluster identity. Of these, 88,294 clusters contained at least one dynamic contact. Sequences within each cluster were aligned to the consensus sequence, and corresponding 3D coordinates were used to compute dynamic contacts (see Figure S16 for a full pipeline overview). Figures S17A–B display the cluster size distributions.

#### Molecular dynamics (MD) simulations

MD data were obtained from the mdCATH^38^ and ATLAS^43^ databases. Dynamic contacts in simulations were computed identically to those in experimental clusters, treating each simulation frame as an independent conformation. All temperature conditions were used for mdCATH.

#### Definition of dynamic contacts

Dynamic contacts were defined as residue–residue pairs (based on C*_α_* atoms) for which at least two structures differed across a contact threshold. Specifically, a pair was considered dynamic if the inter-residue distance was <7.5 Å in one conformation and >8.5 Å in another, introducing a 0.5 Å margin to reduce spurious classifications from small fluctuations.

#### Contact occupancy or frequency

For MD-derived data, we compute the contact occupancy (or frequency) for each residue pair as the fraction of frames in which the contact is formed, aggregated across all trajectory replicates. Contacts are defined using C*_α_* distances with a threshold of 8 Å. Unlike the binary dynamic contact definition, no tolerance margin is applied, as occupancy is a continuous quantity and fluctuations around the threshold provide informative signal about contact stability and uncertainty. This measure can be interpreted as an estimate of the marginal probability of contact formation under the simulated thermodynamic conditions.

#### Contact kinetics

To characterize contact dynamics beyond variability and stability, we compute coarse-grained kinetic features from MD trajectories. For each residue pair, binary contact time series are constructed using an 8 ^Å^ C*_α_* distance threshold. From these time series, we estimate (i) the average on-time (contact lifetime), defined as the mean duration a contact remains continuously formed before breaking, and (ii) the average off-time, defined as the mean duration a residue pair remains separated before forming a contact. These quantities are computed by counting transitions between contact and non-contact states across all trajectory replicates and aggregating the total time spent in each state. Specifically, on-times are given by the total time spent in the contact state divided by the number of contact-breaking events, while off-times are given by the total time spent in the non-contact state divided by the number of contact-forming events. These features capture contact lifetimes and formation timescales within the nanosecond regime accessible to the mdCATH simulations. For downstream modeling, timescales are discretized into coarse-grained bins: always on/off (no transitions observed out of the state), 1–10 ns, 10–100 ns, 100–300 ns, *>*300 ns, and undefined (no transition into the state).

#### Data splits

Datasets were split into training, validation, and test sets (90%/5%/5%) using MMseqs2-based clustering.^6,7^ Clusters assigned to different splits shared no more than 20% sequence identity to ensure independence. 80% minimum coverage was used (-c 0.8) for clustering. For ATLAS, we adopted the test set from AlphaFlow.^27^ Full details on the data splits are provided in the GitHub repository.

#### Model comparisons

For model comparisons, we used the publicly available weights from https://github.com/bjing2016/alphaflow and https://github.com/microsoft/bioemu. All MSAs were generated by querying the ColabFold server^66^ using the corresponding script in https://github.com/bjing2016/alphaflow. For the ATLAS test set, the precomputed ensembles provided by the AlphaFlow authors were employed. For BioEmu, 250 structures were sampled per target to match the number used by AlphaFlow. For the mdCATH test set, 250 structures were sampled per target using each model. An identical pipeline was used for runtime benchmark.

#### Molecular dynamics simulations

All MD simulations for SWEET2b were obtained from ref. 49. No additional MD simulations were performed as part of this study.

### Training procedure

#### Overview

All training was performed on a single NVIDIA RTX 4090 GPU. ESMFold weights were kept frozen throughout (see Table S4).

#### Pretraining

Pretraining was performed on the experimental structure clusters. We note that this pretraining is only relevant for the dynamic contact classification task, and other heads were fitted on MD data only. During each epoch, 10,000 samples were drawn from the training set and 1,000 from the validation set. Proteins were randomly cropped to 256 residues, with sampling probabilities weighted by protein length. Figures S17C–D show that the crop size is appropriate, since *>*93% of dynamic contacts are captured. The Adam optimizer^67^ was used with a learning rate of 1×10*^−^*^4^. The focal loss function^68^ was employed with parameters *α* = 0.99 and *γ* = 2, and a batch size of 64. Training was terminated once the validation loss did not improve for 10 consecutive epochs.

#### Finetuning

Training on the mdCATH dataset followed the same procedure as pretraining, except with 1,000 training samples and 100 validation samples per epoch. To reflect the different class balance, *α* in the focal loss function was adjusted to 0.85. For the frequency head, mean squared error (MSE) was employed. For the kinetics head, weighted multiclass cross-entropy was employed. Class weights used during training for the kinetics head are available in the data repository. To avoid fitting the confidence and residual heads to random noise, the training of these heads was enabled after 10 epochs.

#### Ablation experiments

For the auxiliary input ablation, transition layers were removed. For the DCM ablation, the pairwise representation from the final Evoformer block of ESM-Fold was used in place of the DCM output.

### Validation metrics

#### Balanced accuracy

To address class imbalance, we report balanced accuracy, defined as:

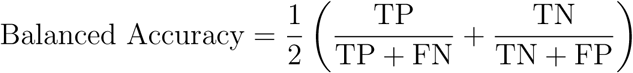

where TP, TN, FP, and FN represent true positives, true negatives, false positives, and false negatives, respectively. This metric reflects the average of recall (sensitivity) and specificity, and treats both classes symmetrically. Other standard classification metrics (precision, recall, F1 score and AUROC), as well as RMSE for regression tasks, are included in Tables S5–23.

#### Resolution metric

To evaluate the spatial precision of dynamic contact classification, we measure the sequence-space distance (L_1_ metric) between predicted and ground-truth positives. For each predicted positive, the minimum L_1_ distance to a true positive is computed. The distance distribution is normalized into a probability density. Faster decay is interpreted as better resolution of dynamic contacts.

## Data availability

Featurized datasets, model weights, and human proteome predictions are available in the Illinois Data Bank at https://doi.org/10.13012/B2IDB-3773897_V2.

## Code availability

All code used to train the model and reproduce the results of the paper is available at https://github.com/ShuklaGroup/esmdynamic.

A user-friendly interface to use the model is available through Google Colab Notebooks at https://colab.research.google.com/github/ShuklaGroup/esmdynamic/blob/main/examples/esmdynamic/esmdynamic.ipynb.

## Acknowledgement

The authors acknowledge support from the National Science Foundation Early CAREER Award (NSF MCB-1845606) and the National Institutes of Health (Award No. R35GM142745). Diego E. Kleiman was supported by a fellowship from The Molecular Sciences Software Institute under NSF grant CHE-2136142.

## Contributions

D.E.K., J.F., and D.S. conceived and designed the study. D.E.K., J.F., and Z.X. collected and analyzed the data. D.E.K. developed the software and machine learning model. D.E.K. drafted the manuscript. J.F., Z.X., and D.S. contributed to manuscript revision and provided feedback. D.S. supervised the study.

## Ethics declarations

### Competing interests

The authors declare no competing interests.

## Supplementary information

Supplementary Methods, Supplementary Results, Tables S1–23, and Figures S1-17.

## Notes

https://github.com/ShuklaGroup/esmdynamic

https://doi.org/10.13012/B2IDB-3773897_V2

### Competing Interest Statement

The authors have declared no competing interest.

### Summary of Updates

Multihead version of ESMDynamic with classification, equilibrium probability regression, kinetics, and confidence modules.

