## Supplementary Information for "ESMDynamic: Fast and Accurate Prediction of Protein Dynamic Contact Maps from Single Sequences"

### 1 Supplementary Methods

#### 1.1 Markov State Modeling

Molecular dynamics trajectories for OsSWEET2b were obtained from ref. S1, comprising a total simulation time of 145 $\mu$s. This dataset is used here in a retrospective setting to evaluate whether collective variables (CVs) derived from sequence-based predictions can recover the same mechanistic features as those obtained from direct trajectory analysis.

**CV selection from ESMDynamic predictions.** The protein sequence was input to ESMDynamic to generate a dynamic contact probability map. Residue–residue distances were selected for featurization following the pipeline shown in Main Text Figure 4A. Briefly, predicted dynamic contacts were filtered to retain only long-range interactions (sequence separation $\geq$ 40 residues), and the resulting set of residue pairs was clustered using single-linkage hierarchical clustering based on $L_1$ distances in contact-map space. The number of clusters was fixed to 9 to match the number of expert-selected CVs reported in ref. S1. From each cluster, the residue pair with the highest predicted dynamic contact probability was selected, and the corresponding inter-residue distances were used as CVs.

**Alternative CV selection protocols.** To enable a controlled comparison, the same CV selection pipeline was applied to two additional sources of contact information: (i) *ESMFold native contacts*, obtained from the predicted structure using an 8 Å $C_\alpha$ distance threshold, and (ii) *MD-derived dynamic contacts*, computed directly from the trajectories using the same definition as in the training data.

For these baselines, no probability scores are available. Therefore, after clustering, a representative pair (i.e., the medoid or geometric center in contact-map space) was selected as the CV for each cluster.

**Feature construction and dimensionality reduction.** For all CV sets, the selected inter-residue distances were computed along the MD trajectories and used to construct feature time series. To reduce dimensionality and extract slow collective motions, time-lagged independent component analysis (tICA)[S2] was applied using the Deeptime implementation.[S3] The 9-dimensional feature space was projected onto the top 3 time-lagged independent components using a lag time consistent with ref. S1.

**Clustering and MSM construction.** The resulting tICA projections were discretized into 900 microstates using mini-batch K-means clustering, following the protocol of ref. S1. A Markov state model (MSM) was then constructed from the discretized trajectories using the maximum-likelihood estimator implemented in Deeptime, with a lag time of 13 ns. This lag time was chosen based on prior work and validated by implied timescale convergence (see Main Text Figure 4D).

**Model validation and comparison.** MSM quality was assessed using the VAMP-2 score,[S4] computed from the top 6 singular values of the Koopman operator to match the methodology of ref. S1. This evaluation was performed consistently across all CV sets, including those derived from ESMDynamic predictions, MD-based dynamic contact maps, and ESMFold native contact maps. All comparisons were conducted using identical clustering and MSM construction protocols.

**Free energy estimation.** MSM-derived stationary probabilities were used to reweight trajectory frames and compute the free energy landscape along the top two tICs using Deeptime.

**Transition path theory.** For the MSM derived with ESMDynamic features, relevant states were identified from the free energy surface. mean-first passage times were computed using Deeptime.

### 2 Supplementary Results

#### 2.1 Supplementary dataset featurization analysis

##### 2.1.1 Dynamic contact comparison in experimental clusters and mdCATH

To assess the consistency of dynamic contact definitions across data sources, we compared dynamic contact maps derived from experimental structure clusters (PDB/AFDB) and MD simulations (mdCATH) for proteins present in both datasets. We find that these definitions are not equivalent and differ substantially in both density and interpretation.

Dynamic contact maps inferred from experimental structural clusters are markedly sparse. This is primarily due to limited sampling, as most clusters contain only a small number of resolved conformations (median of four structures per protein), which restricts the ability to detect contact variability. In addition, experimental structures often rely on stabilization strategies (e.g., crystallization conditions, engineered constructs, or bound partners) that reduce accessible conformational diversity. As a result, only a small fraction of residue pairs are identified as dynamic in these datasets.

In contrast, MD simulations explicitly sample thermal fluctuations within a defined thermodynamic ensemble, enabling frequent formation and disruption of residue–residue contacts over time. This leads to substantially denser dynamic contact maps. Across 3,310 proteins shared between datasets, we observe that $\sim$15% of residue pairs form dynamic contacts in mdCATH simulations at 320 K, compared to only $\sim$0.3% in the corresponding experimental structural clusters.

Beyond differences in density, the two data sources may capture distinct classes of conformational changes. Experimental structures can reflect slower or condition-specific rearrangements (e.g., domain motions, cryptic pocket formation, or ligand-induced conformational changes), whereas MD-derived labels are enriched for faster, local fluctuations within the $\sim$500 ns simulation timescales. Consequently, dynamic contacts derived from these datasets should be interpreted as complementary rather than interchangeable.

Finally, we note that the sparsity of experimentally derived dynamic contact maps has important implications for evaluation. In particular, it leads to regimes characterized by very high recall and very low precision when comparing predicted contacts to experimental labels. This applies both for ESMDynamic as well as BioEmu (see Supplementary Results section 2.3). For this reason, experimental structural clusters are used only during pretraining to expose the model to diverse conformations, while final training objectives and evaluations are based exclusively on MD-derived labels, which provide a more consistent and systematically sampled representation of conformational variability.

A complete correspondence table mapping mdCATH proteins to their associated experimental structural clusters is provided in the data repository. This resource enables direct cross-referencing between datasets and supports reproducibility of the analyses presented in this section.

###### 2.1.2 Dynamic contacts comparison in mdCATH and ATLAS

The mdCATH and ATLAS datasets differ in their sampling frequency, with mdCATH trajectories saved every 1 ns and ATLAS trajectories saved every 10 ps. To evaluate whether this discrepancy affects the assignment of dynamic contacts, we performed a controlled analysis by subsampling the ATLAS trajectories to a 1 ns interval, matching the mdCATH sampling rate, and recomputing the corresponding dynamic contact maps.

We then compared dynamic contact assignments obtained from the original (10 ps) and subsampled (1 ns) ATLAS trajectories. Two metrics were considered: (i) the fraction of residue pairs whose dynamic contact labels differed between sampling rates, and (ii) the absolute mean difference in contact occupancy (i.e., the fraction of frames in which a contact is formed). In both cases, we observe minimal differences. On average, 1.94% of residue pairs changed dynamic-contact labels after subsampling, with a range from 0.34% to 6.71% across proteins. Similarly, the absolute mean difference in contact occupancy was 0.036% on average, with values ranging from 0.0083% to 0.12%.

These results indicate that dynamic contact assignments are largely robust to differences in sampling frequency between mdCATH and ATLAS. Although higher-frequency sampling can, in principle, capture very short-lived fluctuations, the definition of dynamic contacts used here (based on transitions across a distance threshold) appears insensitive to these differences within the range of sampling intervals considered. As a result, dynamic contact labels derived from these datasets can be compared directly without introducing significant bias due to sampling frequency.

Despite this robustness, the two datasets differ substantially in the overall prevalence of dynamic contacts. ATLAS (sampled at 300 K for 100 ns) contains fewer dynamic contacts, with approximately 4.9% of residue pairs classified as dynamic, whereas mdCATH simulations exhibit a higher fraction (15.6% at 320 K, increasing with temperature). This difference reflects reduced conformational variability in ATLAS, resulting in sparser contact patterns. Consequently, the prediction task in ATLAS is biased in favor of recall, which explains the higher balanced accuracy observed for both ESMDynamic and baseline methods (e.g., native contacts achieving ∼0.7 balanced accuracy).

We note, moreover, that the two datasets differ in total trajectory length, with mdCATH simulations extending to 464 ns on average, compared to 100 ns for ATLAS. This difference limits the ability to directly compare kinetic properties across datasets and therefore we refrain from using ATLAS as an external validation set for the kinetics head of ESMDynamic. In particular, shorter ATLAS trajectories may not capture slower contact formation and breaking events, leading to truncated estimates of contact lifetimes.

###### 2.1.3 Analysis of equilibrium convergence in mdCATH trajectories

The finite length of MD trajectories in mdCATH (average 476 ns) limits the extent to which fully converged equilibrium ensembles can be obtained, and successive frames are not strictly independent samples. This is an inherent limitation of MD-based datasets and may impact the estimation of equilibrium properties.

However, the observables used in this work are based on residue-residue contact fluctuations, which are local structural features and may converge more rapidly than global conformational distributions. To assess the robustness of these quantities with respect to sampling, we performed two complementary convergence analyses.

First, we conducted a leave-one-out analysis across trajectory replicates. For each system, dynamic contact maps were constructed using 4 out of the 5 available trajectories, generating five partial-data maps per protein. These were compared to the map obtained using all trajectories, and agreement in dynamic contact assignments was quantified. Second, we performed a time block analysis by splitting each trajectory into two equal halves along the time axis and computing dynamic contact maps independently for each segment. Agreement between these maps was used to assess temporal convergence.

In addition to binary contact labels, we evaluated convergence of contact occupancy (i.e., the fraction of time a contact remains formed). For this quantity, we report the mean absolute error between estimates obtained from the different subsets described above. Across all temperature conditions, both analyses show high agreement in dynamic contact assignments and small deviations in contact occupancy (Table S2), indicating that these observables are largely stable with respect to additional sampling within the timescales probed by mdCATH.

We emphasize that this does not imply full convergence of the underlying conformational ensembles. Longer simulations may reveal additional slow transitions or rare states that are not captured in the current dataset. Rather, these results indicate that the specific features used in this work are locally converged and sufficiently robust for model training and evaluation under the available sampling conditions.

###### 2.1.4 Sensitivity of dynamic contacts to distance cutoff

Dynamic contact assignments are inherently sensitive to the choice of distance cutoff used to define residue-residue interactions. To quantify this effect, we evaluated alternative $C_\alpha$ distance thresholds of 6 Å and 10 Å and compared the resulting dynamic contact maps to those obtained with the default 8 Å threshold using the mdCATH test set (270 proteins). We observe substantial changes in contact assignment, with up to 19% of residue pairs switching classification depending on the cutoff and temperature (Table S3).

In addition, Spearman correlations computed for contact occupancy (fraction of frames in which a contact is formed), as well as for on- and off-times, indicate that featurizations derived from different cutoffs can differ markedly (Table S3). These results highlight that the choice of cutoff directly influences both the sparsity and the information content of the resulting contact maps.

Our choice of an 8 Å $C_\alpha$ distance threshold (with a ±0.5 Å margin) is motivated by both prior literature and empirical considerations. Previous work by Yuan *et al.*[S5] systematically evaluated a wide range of contact definitions and found that $C_\alpha$ cutoffs in the 7–8 Å range provide near-optimal discrimination of protein folds, whereas smaller and larger thresholds degrade performance. In particular, smaller cutoffs (e.g., ≤6 Å) produce overly sparse contact maps that miss relevant interactions, while larger cutoffs (e.g., ≥10 Å) introduce non-specific contacts that reduce structural specificity.

Our definition is consistent with these findings. Due to the ±0.5 Å margin, contacts are effectively labeled as "on" when the $C_\alpha$ distance is below 7.5 Å, aligning with the optimal regime identified in prior studies. The use of a tolerance band around the cutoff reduces sensitivity to small fluctuations arising from numerical noise or thermal motion, ensuring that dynamic contacts reflect meaningful transitions between contact and non-contact states rather than minimal oscillations around a hard threshold.

We additionally adopt a $C_\alpha$-based definition to ensure consistency across all residue types, which is particularly important when comparing contacts across sequence-aligned positions in structural clusters. Taken together, these considerations support the use of an 8 Å $C_\alpha$ cutoff with a tolerance margin as a balanced and robust definition of dynamic contacts, despite the inherent sensitivity of contact assignments to this parameter.

##### 2.1.5 Correlation of dynamic contacts to physicochemical properties

To characterize the structural and physicochemical determinants of dynamic contacts, we analyzed how the probability of observing a dynamic contact depends on properties of the participating residue pairs. Specifically, we computed conditional probabilities of dynamic contact formation as a function of (i) relative solvent accessibility (rSASA), (ii) secondary structure (DSSP classification), and (iii) amino acid type (Supplementary Figures S3–5).

We observe a strong dependence on solvent accessibility. Residue pairs that are both highly solvent-exposed exhibit the highest probability of forming dynamic contacts (exceeding 0.6 in the most exposed regime), whereas buried residue pairs show substantially lower probabilities (∼0.05–0.1). This dependence is monotonic, with dynamic-contact probability increasing as either residue becomes more exposed. This trend is consistent with the expectation that surface residues are more flexible and capable of rearranging interactions, while core residues are constrained by packing.

Dynamic contacts are also strongly modulated by secondary structure. Residues in loop regions show the highest propensity for dynamic contacts, with the presence of a loop residue increasing the likelihood of contact variability regardless of the partner residue. In contrast, ordered secondary structures such as $\alpha$-helices and $\beta$-ladders exhibit lower probabilities (∼0.04–0.09), reflecting their structural rigidity. Notably, rare or irregular motifs such as single $\beta$-bridges and $\pi$-helices display elevated dynamic-contact probabilities, consistent with their known instability and structural flexibility.

In comparison, the dependence on amino acid identity is relatively modest. We observe slightly higher probabilities for polar and structurally atypical residues (e.g., Gly, Pro), and somewhat lower probabilities for certain aromatic or charged pairs. However, the overall variation across residue types is small (∼0.06–0.10), indicating that dynamic contact formation is driven primarily by structural context rather than residue identity alone.

Taken together, these results demonstrate that dynamic contacts are not uniformly distributed across proteins, but are enriched in solvent-exposed, flexible regions and depleted in buried, ordered structural elements. These trends are consistent with the physical picture of protein dynamics captured by MD simulations and support the interpretation of dynamic contacts as markers of fast-motion, local conformational changes.

#### 2.2 Performance comparison on BioEmu benchmarks

To assess whether dynamic-contact maps capture experimentally observed conformational changes, we evaluated ESMDynamic and BioEmu on four benchmark datasets introduced by Lewis et al.:[S6] domain motions, local unfolding, cryptic pocket formation, and OOD60. These datasets were originally designed to evaluate generative models of conformational ensembles; here, we repurpose them for the dynamic contact classification task.

For the domain motion, cryptic pocket, and OOD60 benchmarks, each protein is associated with two experimentally resolved structures representing distinct conformational states. Reference dynamic-contact maps are constructed by identifying residue pairs whose contact state differs between these structures. As discussed in Section 2.1.1, these experimentally derived maps are inherently sparse, leading to evaluation regimes characterized by high recall and low precision for both models. For the local unfolding benchmark, only a single structure is available with annotated unfolding regions; in this case, we define dynamic contacts as native contacts involving these residues and evaluate models based on their ability to recover these disrupted interactions.

A summary of performance across all benchmarks is shown in Table S22. ESMDynamic consistently achieves higher balanced accuracy and recall across all datasets. For example, in OOD60, ESMDynamic attains a balanced accuracy of 0.854 compared to 0.795 for BioEmu, along with substantially higher recall (0.898 vs 0.775). Similar trends are observed in the domain motion and cryptic pocket benchmarks, where ESMDynamic maintains higher balanced accuracy and recall, indicating greater sensitivity to conformational changes. This behavior is also reflected in the local unfolding benchmark, where ESMDynamic achieves slightly higher recall (0.940 vs 0.931).

In contrast, BioEmu consistently achieves higher precision and F1 scores. Across all benchmarks, BioEmu predicts a smaller set of dynamic contacts with higher confidence (e.g., precision of 0.062 vs 0.024 in OOD60), resulting in improved F1 scores despite lower recall. This indicates that BioEmu operates in a more conservative regime, prioritizing specificity over sensitivity.

These results highlight a systematic trade-off between sensitivity and specificity. ESM-Dynamic favors recall, identifying a broader set of potentially dynamic contacts, whereas BioEmu favors precision, identifying fewer but higher-confidence interactions. The higher balanced accuracy of ESMDynamic suggests that, despite lower precision, its predictions better capture the overall distribution of dynamic versus static contacts.

We emphasize that the low precision observed for both models is largely a consequence of label sparsity. Because these experimental benchmarks contain only two resolved conformations, many residue pairs that are dynamic in reality may not be labeled as such, leading to an underestimation of true positives.

Taken together, these results demonstrate that dynamic contact maps (both derived from experimental structures and predicted by ESMDynamic) capture multiple classes of conformational change, including domain rearrangements, local unfolding, and cryptic pocket formation. At the same time, the limitations of experimentally derived labels should be considered when interpreting these benchmarks, which are best viewed as qualitative indicators of sensitivity to conformational change rather than exhaustive measures of predictive accuracy.

#### 2.3 Correlation of dynamic contact predictions to RelaxDB tokens

To relate dynamic contact predictions to experimentally characterized residue-level dynamics, we analyzed their dependence on RelaxDB annotations, which categorize residues according to the timescale of motion measured by NMR. Specifically, RelaxDB assigns discrete tokens to residues indicating distinct dynamical regimes, including fast (ps–ns) motions, slow ($\mu$s–ms) conformational exchange, combined fast and slow dynamics, and disordered or unassigned regions. This provides a natural framework for assessing whether dynamic contacts are associated with particular timescales of motion.

We computed the conditional probability of a residue pair being predicted as a dynamic contact (at $T = 320$ K) as a function of the RelaxDB tokens of the participating residues (Figure S6). We observe a clear enrichment of predicted dynamic contacts among residues annotated with disordered termini ($t$), an intermediate enrichment for residues annotated with fast (ps-ns) motions ($v$) or slow ($\mu$s-ms) exchange ($\hat{}$), and a depletion among residues labeled with both fast and slow motions ($b$) or lacking assignment (.). This trend is consistent with the nature of the MD data used for training: with trajectory lengths of $\sim$500 ns, the dataset predominantly samples fast, local fluctuations, while slower conformational processes are less represented. As a result, ESMDynamic is biased toward detecting contacts associated with short-timescale dynamics.

Taken together, these results indicate that dynamic contacts predicted by ESMDynamic are primarily associated with fast, local motions, but nevertheless show meaningful correlations with experimentally derived residue-level dynamics. This analysis helps clarify the interpretation of "dynamic contacts" in our framework: rather than capturing all potential timescales, they predominantly reflect interactions modulated by fast fluctuations within the range accessible to the MD datasets.

#### 2.4 Analysis of predictions for intrinsically disordered proteins

Intrinsically disordered proteins (IDPs) represent a challenging regime for models of protein dynamics, as they lack a well-defined folded structure and instead populate broad conformational ensembles. In such systems, residue–residue interactions are inherently transient, and dynamic contacts are expected to play a dominant role in describing their behavior. To assess the performance of ESMDynamic in this regime, we analyzed representative IDPs, specifically $\alpha$-synuclein and tau, using available experimental structural clusters.

For these proteins, we constructed reference dynamic contact maps from experimental structure clusters (clusters #405 and #607, involving PDB IDs 1XQ8 and 6HRE, for reference). These maps were compared against predictions from ESMDynamic and native contact maps derived from ESMFold (Figure S15). Unlike the MD-based datasets used throughout the Main Text, these experimental clusters are highly sparse and reflect limited sampling of conformational variability, which must be taken into account when interpreting results.

We observe that ESMDynamic performance is substantially degraded on these IDP systems, in contrast to its strong performance on structured proteins. In particular, ESMDynamic predictions fall below the ESMFold native-contact baseline, indicating that the model struggles to accurately identify dynamic contacts in highly disordered sequences. This behavior is consistent across both $\alpha$-synuclein and tau protein, suggesting a systematic limitation rather than a protein-specific effect. In $\alpha$-synuclein, ESMDynamic obtains 60% balanced accuracy, while the ESMFold baseline obtains 68%. On the tau protein, ESMDynamic obtains 57%, while ESMFold obtains 59%.

This degradation can be explained by two primary factors. First, the mdCATH dataset used for fine-tuning consists predominantly of ordered, globular proteins, and therefore the model is optimized for detecting contact variability within a well-defined structural scaffold. In contrast, IDPs lack persistent tertiary structure, leading to fundamentally different patterns of residue interactions that are not well represented in the training data. Second, ESMDynamic predictions are strongly correlated with the confidence of the underlying structure prediction model (pLDDT), which is typically low for IDPs. As a result, both the structural prior and the learned dynamic features become less reliable in disordered regions.

In summary, while ESMDynamic provides meaningful predictions for structured proteins and systems with well-defined conformational transitions, caution should be exercised when applying it to IDPs, where both the structural inputs and the learned representations are less informative. We include these results in Figure S15 to provide a complete and transparent assessment of the model's scope and limitations.

### 3 Supplementary Tables

Table S1: Overlap between dynamic and native contacts across temperatures in mdCATH dataset. Values are reported as mean ± standard deviation (%).
Dynamic also native = $|\text{dynamic} \cap \text{native}| / |\text{dynamic}|$
Native also dynamic = $|\text{dynamic} \cap \text{native}| / |\text{native}|$

| T (K) | Dynamic also native | Native also dynamic |
|---|---|---|
| 320 | 27.3 ± 10.8 | 64.6 ± 14.6 |
| 348 | 22.4 ± 9.8 | 72.7 ± 14.4 |
| 379 | 16.9 ± 8.4 | 82.0 ± 13.1 |
| 413 | 11.2 ± 6.4 | 91.4 ± 10.1 |
| 450 | 6.8 ± 3.6 | 97.8 ± 5.2 |

Table S2: Summary of label agreement and occupancy MAE across temperatures for leave-one-out and temporal block analyses. Values are reported as mean ± standard deviation.

| | Leave-one-out | | Time Block | |
|---|---|---|---|---|
| T (K) | Label agreement (%) | Occupancy MAE (%) | Label agreement (%) | Occupancy MAE (%) |
| 320 | $98.95 \pm 0.68$ | $0.213 \pm 0.042$ | $97.21 \pm 0.87$ | $0.365 \pm 1.28 \times 10^{-6}$ |
| 348 | $98.65 \pm 1.00$ | $0.248 \pm 0.049$ | $96.29 \pm 1.46$ | $0.457 \pm 2.33 \times 10^{-6}$ |
| 379 | $98.24 \pm 1.47$ | $0.287 \pm 0.057$ | $94.79 \pm 2.69$ | $0.598 \pm 2.80 \times 10^{-6}$ |
| 413 | $97.76 \pm 2.07$ | $0.316 \pm 0.063$ | $92.49 \pm 5.03$ | $0.794 \pm 4.13 \times 10^{-6}$ |
| 450 | $97.84 \pm 1.84$ | $0.294 \pm 0.056$ | $91.21 \pm 6.41$ | $0.918 \pm 4.71 \times 10^{-6}$ |

Table S3: Effect of distance cutoff on agreement and correlation metrics across temperatures (mdCATH test set). Values are reported as mean ± standard deviation (%).

| Cutoff | T (K) | Label agreement | $\rho$ occupancy | $\rho$ on-time | $\rho$ off-time |
|---|---|---|---|---|---|
| 6 Å | 320 | $89.95 \pm 7.69$ | $74.80 \pm 4.36$ | $53.49 \pm 9.36$ | $41.35 \pm 9.60$ |
| | 348 | $88.37 \pm 7.71$ | $76.90 \pm 5.10$ | $60.08 \pm 7.59$ | $47.33 \pm 7.95$ |
| | 379 | $86.03 \pm 8.15$ | $79.82 \pm 5.28$ | $64.97 \pm 5.90$ | $51.80 \pm 6.08$ |
| | 413 | $83.73 \pm 8.21$ | $83.54 \pm 4.87$ | $69.95 \pm 4.42$ | $56.59 \pm 4.80$ |
| | 450 | $81.44 \pm 8.81$ | $86.79 \pm 3.31$ | $72.72 \pm 4.89$ | $59.48 \pm 3.65$ |
| 10 Å | 320 | $86.91 \pm 5.36$ | $77.46 \pm 5.14$ | $48.85 \pm 10.33$ | $46.19 \pm 11.32$ |
| | 348 | $86.63 \pm 5.36$ | $80.03 \pm 5.45$ | $56.50 \pm 10.27$ | $51.65 \pm 10.35$ |
| | 379 | $87.00 \pm 5.76$ | $83.36 \pm 5.45$ | $64.55 \pm 9.93$ | $57.81 \pm 8.82$ |
| | 413 | $88.94 \pm 6.58$ | $87.54 \pm 5.02$ | $74.51 \pm 9.14$ | $64.07 \pm 7.55$ |
| | 450 | $92.80 \pm 6.79$ | $91.76 \pm 3.62$ | $83.33 \pm 6.85$ | $69.26 \pm 5.39$ |

Table S4: ESMDynamic submodules and number of parameters.

| **Module** | **# params** | **Trainable** |
|---|---|---|
| ESMFold | 3,531,600,228 | No |
| Sequence transition (3X) | 2,972,322 | Yes |
| Pair transition (3X) | 33,280 | Yes |
| Dynamic Contact Module (3X) | 28,600,448 | Yes |
| Pairwise position embedding | 8,448 | |
| Evoformer (2X) | 14,293,888 | |
| Layer norm. | 2,048 | |
| Sequence to pair (lin. proj.) | 149,760 | |
| Pair to sequence (lin. proj.) | 4,352 | |
| Sequence attn. | 5,244,928 | |
| Triangle mult. update (outgoing) | 99,584 | |
| Triangle mult. update (incoming) | 99,584 | |
| Triangular attn. (starting) | 82,944 | |
| Triangular attn. (ending) | 82,944 | |
| MLP sequence | 8,395,776 | |
| MLP pair | 131,968 | |
| Recycle s norm | 2,048 | |
| Recycle z norm | 256 | |
| Recycle disto | 1,920 | |
| Prediction layer (classification) | 645 | Yes |
| Confidence head (classification) | 529,413 | Yes |
| Prediction layer (frequency) | 645 | Yes |
| Residual head (frequency) | 8,837 | Yes |
| Prediction layer (kinetics) | 7,740 | Yes |
| Confidence head (kinetics) | 529,413 | Yes |
| Total | 3,627,495,071 | 95,894,843 (2.6%) |

Table S5: Dynamic contact classification head performance. Values are reported as mean ± standard error.

| Dataset | Metric | 320 K | 348 K | 379 K | 413 K | 450 K |
|---|---|---|---|---|---|---|
| Training | balanced accuracy | 0.811 ± 0.002 | 0.786 ± 0.002 | 0.742 ± 0.002 | 0.654 ± 0.002 | 0.554 ± 0.002 |
| | precision | 0.496 ± 0.003 | 0.515 ± 0.003 | 0.555 ± 0.003 | 0.632 ± 0.004 | 0.790 ± 0.004 |
| | recall | 0.792 ± 0.002 | 0.830 ± 0.002 | 0.858 ± 0.002 | 0.920 ± 0.001 | 0.993 ± 0.000 |
| | F1 score | 0.574 ± 0.002 | 0.597 ± 0.002 | 0.633 ± 0.002 | 0.711 ± 0.003 | 0.851 ± 0.003 |
| | AUROC | 0.900 ± 0.001 | 0.881 ± 0.002 | 0.850 ± 0.002 | 0.797 ± 0.002 | 0.722 ± 0.002 |
| Validation | balanced accuracy | 0.812 ± 0.006 | 0.793 ± 0.007 | 0.739 ± 0.008 | 0.667 ± 0.009 | 0.557 ± 0.009 |
| | precision | 0.498 ± 0.011 | 0.513 ± 0.012 | 0.553 ± 0.014 | 0.622 ± 0.016 | 0.784 ± 0.016 |
| | recall | 0.792 ± 0.009 | 0.841 ± 0.008 | 0.860 ± 0.008 | 0.928 ± 0.006 | 0.996 ± 0.001 |
| | F1 score | 0.574 ± 0.007 | 0.599 ± 0.009 | 0.630 ± 0.010 | 0.707 ± 0.013 | 0.848 ± 0.012 |
| | AUROC | 0.900 ± 0.006 | 0.886 ± 0.006 | 0.853 ± 0.007 | 0.810 ± 0.007 | 0.735 ± 0.008 |
| Test | balanced accuracy | 0.796 ± 0.007 | 0.772 ± 0.007 | 0.734 ± 0.008 | 0.657 ± 0.008 | 0.542 ± 0.007 |
| | precision | 0.511 ± 0.012 | 0.533 ± 0.013 | 0.576 ± 0.014 | 0.646 ± 0.016 | 0.791 ± 0.016 |
| | recall | 0.767 ± 0.010 | 0.809 ± 0.009 | 0.844 ± 0.009 | 0.918 ± 0.006 | 0.994 ± 0.001 |
| | F1 score | 0.569 ± 0.008 | 0.596 ± 0.009 | 0.639 ± 0.010 | 0.720 ± 0.012 | 0.853 ± 0.012 |
| | AUROC | 0.889 ± 0.006 | 0.873 ± 0.007 | 0.841 ± 0.008 | 0.796 ± 0.008 | 0.729 ± 0.007 |

Table S6: Contact frequency regression head performance. Values are reported as mean ± standard error.

| Dataset | Metric | 320 K | 348 K | 379 K | 413 K | 450 K |
|---|---|---|---|---|---|---|
| Training | RMSE | 0.073 ± 0.001 | 0.071 ± 0.000 | 0.070 ± 0.000 | 0.066 ± 0.000 | 0.057 ± 0.000 |
| Validation | RMSE | 0.071 ± 0.002 | 0.070 ± 0.002 | 0.068 ± 0.001 | 0.065 ± 0.001 | 0.055 ± 0.001 |
| Test | RMSE | 0.076 ± 0.002 | 0.074 ± 0.002 | 0.072 ± 0.002 | 0.068 ± 0.002 | 0.057 ± 0.002 |

Table S7: On-time kinetics head performance (macro-averaged metrics). Values are reported as mean ± standard error. Note recall and balanced accuracy are equivalent under macro averaging.

| Dataset | Metric | 320 K | 348 K | 379 K | 413 K | 450 K |
|---|---|---|---|---|---|---|
| | balanced accuracy | 0.343 ± 0.001 | 0.345 ± 0.001 | 0.367 ± 0.001 | 0.418 ± 0.001 | 0.540 ± 0.002 |
| | precision | 0.346 ± 0.001 | 0.328 ± 0.001 | 0.315 ± 0.001 | 0.304 ± 0.001 | 0.302 ± 0.001 |
| Training | recall | 0.343 ± 0.001 | 0.345 ± 0.001 | 0.367 ± 0.001 | 0.418 ± 0.001 | 0.540 ± 0.002 |
| | F1 score | 0.191 ± 0.001 | 0.187 ± 0.001 | 0.203 ± 0.001 | 0.222 ± 0.001 | 0.274 ± 0.001 |
| | AUROC | 0.898 ± 0.001 | 0.893 ± 0.001 | 0.889 ± 0.001 | 0.899 ± 0.001 | 0.908 ± 0.001 |
| | balanced accuracy | 0.341 ± 0.003 | 0.346 ± 0.003 | 0.368 ± 0.003 | 0.424 ± 0.007 | 0.545 ± 0.011 |
| | precision | 0.336 ± 0.005 | 0.325 ± 0.003 | 0.311 ± 0.004 | 0.305 ± 0.004 | 0.310 ± 0.005 |
| Validation | recall | 0.341 ± 0.003 | 0.346 ± 0.003 | 0.368 ± 0.003 | 0.424 ± 0.007 | 0.545 ± 0.011 |
| | F1 score | 0.189 ± 0.003 | 0.185 ± 0.003 | 0.199 ± 0.004 | 0.225 ± 0.004 | 0.283 ± 0.005 |
| | AUROC | 0.902 ± 0.005 | 0.897 ± 0.005 | 0.893 ± 0.004 | 0.898 ± 0.004 | 0.911 ± 0.005 |
| | balanced accuracy | 0.339 ± 0.003 | 0.343 ± 0.003 | 0.367 ± 0.003 | 0.409 ± 0.006 | 0.537 ± 0.010 |
| | precision | 0.340 ± 0.005 | 0.324 ± 0.004 | 0.312 ± 0.004 | 0.301 ± 0.004 | 0.292 ± 0.004 |
| Test | recall | 0.339 ± 0.003 | 0.343 ± 0.003 | 0.367 ± 0.003 | 0.409 ± 0.006 | 0.537 ± 0.010 |
| | F1 score | 0.187 ± 0.003 | 0.181 ± 0.003 | 0.196 ± 0.004 | 0.214 ± 0.004 | 0.267 ± 0.004 |
| | AUROC | 0.891 ± 0.006 | 0.887 ± 0.006 | 0.887 ± 0.005 | 0.892 ± 0.005 | 0.897 ± 0.008 |

Table S8: Off-time kinetics head performance (macro-averaged metrics). Values are reported as mean ± standard error. Note recall and balanced accuracy are equivalent under macro averaging.

| Dataset | Metric | 320 K | 348 K | 379 K | 413 K | 450 K |
|---|---|---|---|---|---|---|
| Training | balanced accuracy | $0.438 \pm 0.001$ | $0.439 \pm 0.001$ | $0.436 \pm 0.001$ | $0.439 \pm 0.001$ | $0.457 \pm 0.001$ |
| | precision | $0.483 \pm 0.001$ | $0.470 \pm 0.001$ | $0.444 \pm 0.001$ | $0.407 \pm 0.001$ | $0.412 \pm 0.001$ |
| | recall | $0.438 \pm 0.001$ | $0.439 \pm 0.001$ | $0.436 \pm 0.001$ | $0.439 \pm 0.001$ | $0.457 \pm 0.001$ |
| | F1 score | $0.377 \pm 0.001$ | $0.370 \pm 0.001$ | $0.354 \pm 0.001$ | $0.342 \pm 0.001$ | $0.370 \pm 0.001$ |
| | AUROC | $0.894 \pm 0.001$ | $0.882 \pm 0.001$ | $0.862 \pm 0.001$ | $0.834 \pm 0.001$ | $0.797 \pm 0.001$ |
| Validation | balanced accuracy | $0.436 \pm 0.003$ | $0.441 \pm 0.003$ | $0.436 \pm 0.003$ | $0.442 \pm 0.004$ | $0.465 \pm 0.004$ |
| | precision | $0.474 \pm 0.005$ | $0.469 \pm 0.005$ | $0.444 \pm 0.006$ | $0.412 \pm 0.005$ | $0.426 \pm 0.005$ |
| | recall | $0.436 \pm 0.003$ | $0.441 \pm 0.003$ | $0.436 \pm 0.003$ | $0.442 \pm 0.004$ | $0.465 \pm 0.004$ |
| | F1 score | $0.370 \pm 0.005$ | $0.369 \pm 0.005$ | $0.350 \pm 0.005$ | $0.345 \pm 0.004$ | $0.382 \pm 0.005$ |
| | AUROC | $0.896 \pm 0.005$ | $0.887 \pm 0.005$ | $0.866 \pm 0.004$ | $0.834 \pm 0.005$ | $0.798 \pm 0.005$ |
| Test | balanced accuracy | $0.432 \pm 0.004$ | $0.433 \pm 0.004$ | $0.430 \pm 0.004$ | $0.434 \pm 0.004$ | $0.451 \pm 0.004$ |
| | precision | $0.471 \pm 0.006$ | $0.458 \pm 0.006$ | $0.430 \pm 0.007$ | $0.402 \pm 0.006$ | $0.410 \pm 0.005$ |
| | recall | $0.432 \pm 0.004$ | $0.433 \pm 0.004$ | $0.430 \pm 0.004$ | $0.434 \pm 0.004$ | $0.451 \pm 0.004$ |
| | F1 score | $0.365 \pm 0.005$ | $0.358 \pm 0.006$ | $0.343 \pm 0.006$ | $0.334 \pm 0.005$ | $0.369 \pm 0.004$ |
| | AUROC | $0.884 \pm 0.006$ | $0.872 \pm 0.006$ | $0.854 \pm 0.006$ | $0.828 \pm 0.006$ | $0.796 \pm 0.005$ |

Table S9: Dynamic contact confidence head performance. Values are reported as mean ± standard error.

| Dataset | Metric | 320 K | 348 K | 379 K | 413 K | 450 K |
|---|---|---|---|---|---|---|
| Training | RMSE | 0.080 ± 0.001 | 0.104 ± 0.001 | 0.115 ± 0.001 | 0.163 ± 0.001 | 0.175 ± 0.001 |
| Validation | RMSE | 0.080 ± 0.003 | 0.102 ± 0.004 | 0.117 ± 0.004 | 0.162 ± 0.005 | 0.173 ± 0.006 |
| Test | RMSE | 0.087 ± 0.004 | 0.109 ± 0.005 | 0.123 ± 0.005 | 0.170 ± 0.006 | 0.180 ± 0.006 |

Table S10: Contact frequency residual head performance. Values are reported as mean ± standard error.

| Dataset | Metric | 320 K | 348 K | 379 K | 413 K | 450 K |
|---|---|---|---|---|---|---|
| Training | RMSE | 0.053 ± 0.000 | 0.050 ± 0.000 | 0.048 ± 0.000 | 0.043 ± 0.000 | 0.040 ± 0.000 |
| Validation | RMSE | 0.052 ± 0.002 | 0.049 ± 0.001 | 0.046 ± 0.001 | 0.042 ± 0.001 | 0.038 ± 0.001 |
| Test | RMSE | 0.056 ± 0.002 | 0.053 ± 0.002 | 0.050 ± 0.002 | 0.045 ± 0.001 | 0.040 ± 0.001 |

Table S11: Kinetics confidence head performance. Values are reported as mean ± standard error.

| Dataset | Metric | 320 K | 348 K | 379 K | 413 K | 450 K |
|---|---|---|---|---|---|---|
| Training | RMSE | 0.096 ± 0.001 | 0.115 ± 0.001 | 0.136 ± 0.001 | 0.160 ± 0.001 | 0.131 ± 0.001 |
| Validation | RMSE | 0.098 ± 0.004 | 0.116 ± 0.004 | 0.140 ± 0.005 | 0.161 ± 0.004 | 0.129 ± 0.003 |
| Test | RMSE | 0.102 ± 0.005 | 0.120 ± 0.005 | 0.142 ± 0.005 | 0.159 ± 0.004 | 0.133 ± 0.004 |

Table S12: Dynamic contact classification head performance for the ablated model without auxiliary inputs. Values are reported as mean ± standard error.

| Dataset | Metric | 320 K | 348 K | 379 K | 413 K | 450 K |
|---|---|---|---|---|---|---|
| | balanced accuracy | 0.786 ± 0.001 | 0.789 ± 0.002 | 0.699 ± 0.002 | 0.638 ± 0.002 | 0.559 ± 0.002 |
| | precision | 0.453 ± 0.002 | 0.500 ± 0.003 | 0.494 ± 0.003 | 0.610 ± 0.004 | 0.792 ± 0.004 |
| Training | recall | 0.751 ± 0.002 | 0.803 ± 0.002 | 0.816 ± 0.002 | 0.936 ± 0.001 | 0.964 ± 0.001 |
| | F1 score | 0.531 ± 0.002 | 0.578 ± 0.002 | 0.569 ± 0.002 | 0.698 ± 0.003 | 0.844 ± 0.003 |
| | AUROC | 0.871 ± 0.001 | 0.872 ± 0.002 | 0.797 ± 0.002 | 0.783 ± 0.002 | 0.655 ± 0.002 |
| | balanced accuracy | 0.787 ± 0.006 | 0.795 ± 0.006 | 0.694 ± 0.007 | 0.649 ± 0.009 | 0.563 ± 0.008 |
| | precision | 0.456 ± 0.011 | 0.500 ± 0.012 | 0.496 ± 0.015 | 0.597 ± 0.017 | 0.787 ± 0.016 |
| Validation | recall | 0.753 ± 0.009 | 0.812 ± 0.008 | 0.816 ± 0.008 | 0.944 ± 0.005 | 0.968 ± 0.003 |
| | F1 score | 0.532 ± 0.007 | 0.580 ± 0.008 | 0.568 ± 0.010 | 0.691 ± 0.014 | 0.842 ± 0.012 |
| | AUROC | 0.870 ± 0.006 | 0.877 ± 0.006 | 0.793 ± 0.008 | 0.796 ± 0.007 | 0.669 ± 0.008 |
| | balanced accuracy | 0.771 ± 0.006 | 0.778 ± 0.006 | 0.690 ± 0.008 | 0.639 ± 0.008 | 0.548 ± 0.007 |
| | precision | 0.466 ± 0.012 | 0.517 ± 0.013 | 0.516 ± 0.015 | 0.623 ± 0.017 | 0.794 ± 0.016 |
| Test | recall | 0.731 ± 0.009 | 0.779 ± 0.009 | 0.806 ± 0.009 | 0.935 ± 0.005 | 0.966 ± 0.003 |
| | F1 score | 0.526 ± 0.008 | 0.576 ± 0.008 | 0.580 ± 0.010 | 0.709 ± 0.013 | 0.846 ± 0.012 |
| | AUROC | 0.860 ± 0.007 | 0.864 ± 0.007 | 0.784 ± 0.008 | 0.783 ± 0.008 | 0.655 ± 0.008 |

Table S13: Contact frequency regression head performance for the ablated model without auxiliary inputs. Values are reported as mean ± standard error.

| Dataset | Metric | 320 K | 348 K | 379 K | 413 K | 450 K |
|---|---|---|---|---|---|---|
| Training | RMSE | 0.092 ± 0.001 | 0.083 ± 0.001 | 0.075 ± 0.000 | 0.084 ± 0.000 | 0.070 ± 0.000 |
| Validation | RMSE | 0.091 ± 0.002 | 0.081 ± 0.002 | 0.073 ± 0.002 | 0.083 ± 0.002 | 0.067 ± 0.001 |
| Test | RMSE | 0.095 ± 0.002 | 0.086 ± 0.002 | 0.078 ± 0.002 | 0.085 ± 0.002 | 0.069 ± 0.001 |

Table S14: On-time kinetics classification head performance for the ablated model without auxiliary inputs (macro-averaged metrics). Values are reported as mean ± standard error.

| Dataset | Metric | 320 K | 348 K | 379 K | 413 K | 450 K |
|---|---|---|---|---|---|---|
| Training | balanced accuracy | 0.344 ± 0.001 | 0.350 ± 0.001 | 0.357 ± 0.001 | 0.411 ± 0.001 | 0.497 ± 0.002 |
| | precision | 0.284 ± 0.001 | 0.278 ± 0.001 | 0.270 ± 0.001 | 0.260 ± 0.001 | 0.217 ± 0.001 |
| | recall | 0.344 ± 0.001 | 0.350 ± 0.001 | 0.357 ± 0.001 | 0.411 ± 0.001 | 0.497 ± 0.002 |
| | F1 score | 0.205 ± 0.001 | 0.208 ± 0.001 | 0.201 ± 0.001 | 0.214 ± 0.001 | 0.208 ± 0.001 |
| | AUROC | 0.850 ± 0.001 | 0.825 ± 0.001 | 0.838 ± 0.001 | 0.855 ± 0.001 | 0.824 ± 0.001 |
| Validation | balanced accuracy | 0.342 ± 0.003 | 0.349 ± 0.003 | 0.355 ± 0.004 | 0.414 ± 0.006 | 0.505 ± 0.010 |
| | precision | 0.280 ± 0.003 | 0.273 ± 0.004 | 0.264 ± 0.003 | 0.256 ± 0.003 | 0.219 ± 0.003 |
| | recall | 0.342 ± 0.003 | 0.349 ± 0.003 | 0.355 ± 0.004 | 0.414 ± 0.006 | 0.505 ± 0.010 |
| | F1 score | 0.202 ± 0.003 | 0.205 ± 0.003 | 0.196 ± 0.004 | 0.213 ± 0.004 | 0.211 ± 0.003 |
| | AUROC | 0.854 ± 0.005 | 0.826 ± 0.005 | 0.840 ± 0.006 | 0.849 ± 0.005 | 0.830 ± 0.005 |
| Test | balanced accuracy | 0.339 ± 0.003 | 0.343 ± 0.003 | 0.349 ± 0.004 | 0.406 ± 0.005 | 0.492 ± 0.010 |
| | precision | 0.278 ± 0.003 | 0.272 ± 0.004 | 0.265 ± 0.004 | 0.258 ± 0.004 | 0.215 ± 0.003 |
| | recall | 0.339 ± 0.003 | 0.343 ± 0.003 | 0.349 ± 0.004 | 0.406 ± 0.005 | 0.492 ± 0.010 |
| | F1 score | 0.199 ± 0.003 | 0.202 ± 0.003 | 0.190 ± 0.004 | 0.208 ± 0.004 | 0.207 ± 0.003 |
| | AUROC | 0.842 ± 0.007 | 0.819 ± 0.006 | 0.835 ± 0.007 | 0.848 ± 0.006 | 0.811 ± 0.008 |

Table S15: Off-time kinetics classification head performance for the ablated model without auxiliary inputs (macro-averaged metrics). Values are reported as mean ± standard error.

| Dataset | Metric | 320 K | 348 K | 379 K | 413 K | 450 K |
|---|---|---|---|---|---|---|
| Training | balanced accuracy | 0.414 ± 0.001 | 0.424 ± 0.001 | 0.415 ± 0.001 | 0.421 ± 0.001 | 0.437 ± 0.001 |
| | precision | 0.373 ± 0.001 | 0.367 ± 0.001 | 0.357 ± 0.001 | 0.323 ± 0.001 | 0.312 ± 0.001 |
| | recall | 0.414 ± 0.001 | 0.424 ± 0.001 | 0.415 ± 0.001 | 0.421 ± 0.001 | 0.437 ± 0.001 |
| | F1 score | 0.355 ± 0.001 | 0.354 ± 0.001 | 0.342 ± 0.001 | 0.312 ± 0.001 | 0.305 ± 0.001 |
| | AUROC | 0.842 ± 0.001 | 0.815 ± 0.001 | 0.804 ± 0.001 | 0.789 ± 0.001 | 0.766 ± 0.001 |
| Validation | balanced accuracy | 0.414 ± 0.003 | 0.426 ± 0.004 | 0.415 ± 0.003 | 0.425 ± 0.003 | 0.443 ± 0.004 |
| | precision | 0.372 ± 0.003 | 0.364 ± 0.004 | 0.357 ± 0.004 | 0.321 ± 0.004 | 0.316 ± 0.003 |
| | recall | 0.414 ± 0.003 | 0.426 ± 0.004 | 0.415 ± 0.003 | 0.425 ± 0.003 | 0.443 ± 0.004 |
| | F1 score | 0.352 ± 0.005 | 0.353 ± 0.005 | 0.340 ± 0.004 | 0.314 ± 0.004 | 0.311 ± 0.003 |
| | AUROC | 0.843 ± 0.005 | 0.819 ± 0.005 | 0.806 ± 0.004 | 0.789 ± 0.005 | 0.768 ± 0.005 |
| Test | balanced accuracy | 0.409 ± 0.004 | 0.418 ± 0.004 | 0.410 ± 0.004 | 0.417 ± 0.004 | 0.432 ± 0.004 |
| | precision | 0.363 ± 0.004 | 0.359 ± 0.005 | 0.349 ± 0.004 | 0.320 ± 0.004 | 0.312 ± 0.003 |
| | recall | 0.409 ± 0.004 | 0.418 ± 0.004 | 0.410 ± 0.004 | 0.417 ± 0.004 | 0.432 ± 0.004 |
| | F1 score | 0.345 ± 0.005 | 0.343 ± 0.006 | 0.334 ± 0.005 | 0.308 ± 0.005 | 0.306 ± 0.003 |
| | AUROC | 0.833 ± 0.006 | 0.804 ± 0.006 | 0.797 ± 0.005 | 0.784 ± 0.007 | 0.767 ± 0.006 |

Table S16: Dynamic contact classification head performance for the ablated model without the dynamic contact module (DCM). Values are reported as mean ± standard error.

| Dataset | Metric | 320 K | 348 K | 379 K | 413 K | 450 K |
|---|---|---|---|---|---|---|
| Training | balanced accuracy | 0.775 ± 0.001 | 0.777 ± 0.002 | 0.730 ± 0.002 | 0.630 ± 0.002 | 0.562 ± 0.002 |
| | precision | 0.427 ± 0.002 | 0.499 ± 0.003 | 0.531 ± 0.003 | 0.604 ± 0.004 | 0.792 ± 0.004 |
| | recall | 0.713 ± 0.002 | 0.790 ± 0.002 | 0.825 ± 0.002 | 0.920 ± 0.001 | 0.973 ± 0.001 |
| | F1 score | 0.503 ± 0.001 | 0.575 ± 0.002 | 0.604 ± 0.002 | 0.688 ± 0.003 | 0.846 ± 0.003 |
| | AUROC | 0.848 ± 0.001 | 0.864 ± 0.002 | 0.825 ± 0.002 | 0.759 ± 0.002 | 0.667 ± 0.002 |
| Validation | balanced accuracy | 0.774 ± 0.006 | 0.783 ± 0.006 | 0.729 ± 0.007 | 0.644 ± 0.009 | 0.563 ± 0.008 |
| | precision | 0.429 ± 0.011 | 0.499 ± 0.012 | 0.531 ± 0.015 | 0.597 ± 0.017 | 0.787 ± 0.016 |
| | recall | 0.716 ± 0.008 | 0.802 ± 0.008 | 0.831 ± 0.008 | 0.920 ± 0.005 | 0.974 ± 0.002 |
| | F1 score | 0.502 ± 0.006 | 0.578 ± 0.008 | 0.605 ± 0.010 | 0.684 ± 0.013 | 0.843 ± 0.012 |
| | AUROC | 0.847 ± 0.006 | 0.869 ± 0.006 | 0.824 ± 0.007 | 0.771 ± 0.007 | 0.674 ± 0.008 |
| Test | balanced accuracy | 0.764 ± 0.006 | 0.763 ± 0.007 | 0.720 ± 0.008 | 0.631 ± 0.008 | 0.550 ± 0.007 |
| | precision | 0.439 ± 0.011 | 0.514 ± 0.013 | 0.549 ± 0.014 | 0.617 ± 0.017 | 0.794 ± 0.016 |
| | recall | 0.695 ± 0.008 | 0.770 ± 0.009 | 0.813 ± 0.009 | 0.915 ± 0.005 | 0.972 ± 0.003 |
| | F1 score | 0.501 ± 0.007 | 0.571 ± 0.008 | 0.611 ± 0.010 | 0.697 ± 0.013 | 0.847 ± 0.012 |
| | AUROC | 0.839 ± 0.006 | 0.855 ± 0.007 | 0.815 ± 0.007 | 0.753 ± 0.008 | 0.671 ± 0.006 |

Table S17: Contact frequency regression head performance for the ablated model without the dynamic contact module (DCM). Values are reported as mean ± standard error.

| Dataset | Metric | 320 K | 348 K | 379 K | 413 K | 450 K |
|---|---|---|---|---|---|---|
| Training | RMSE | 0.187 ± 0.001 | 0.100 ± 0.000 | 0.110 ± 0.000 | 0.108 ± 0.000 | 0.100 ± 0.000 |
| Validation | RMSE | 0.184 ± 0.003 | 0.099 ± 0.002 | 0.108 ± 0.002 | 0.107 ± 0.002 | 0.098 ± 0.002 |
| Test | RMSE | 0.183 ± 0.003 | 0.102 ± 0.002 | 0.111 ± 0.002 | 0.107 ± 0.002 | 0.099 ± 0.002 |

Table S18: On-time kinetics classification head performance for the ablated model without the dynamic contact module (DCM) (macro-averaged metrics). Values are reported as mean ± standard error.

| Dataset | Metric | 320 K | 348 K | 379 K | 413 K | 450 K |
|---|---|---|---|---|---|---|
| Training | balanced accuracy | 0.353 ± 0.001 | 0.349 ± 0.001 | 0.357 ± 0.001 | 0.397 ± 0.001 | 0.475 ± 0.002 |
| | precision | 0.326 ± 0.001 | 0.319 ± 0.001 | 0.261 ± 0.001 | 0.252 ± 0.001 | 0.215 ± 0.001 |
| | recall | 0.353 ± 0.001 | 0.349 ± 0.001 | 0.357 ± 0.001 | 0.397 ± 0.001 | 0.475 ± 0.002 |
| | F1 score | 0.213 ± 0.001 | 0.202 ± 0.001 | 0.197 ± 0.001 | 0.209 ± 0.001 | 0.205 ± 0.001 |
| | AUROC | 0.870 ± 0.001 | 0.861 ± 0.001 | 0.828 ± 0.001 | 0.859 ± 0.001 | 0.825 ± 0.001 |
| Validation | balanced accuracy | 0.350 ± 0.003 | 0.347 ± 0.003 | 0.359 ± 0.003 | 0.396 ± 0.005 | 0.488 ± 0.010 |
| | precision | 0.317 ± 0.004 | 0.313 ± 0.004 | 0.258 ± 0.003 | 0.250 ± 0.004 | 0.216 ± 0.003 |
| | recall | 0.350 ± 0.003 | 0.347 ± 0.003 | 0.359 ± 0.003 | 0.396 ± 0.005 | 0.488 ± 0.010 |
| | F1 score | 0.208 ± 0.003 | 0.200 ± 0.004 | 0.193 ± 0.004 | 0.211 ± 0.004 | 0.208 ± 0.003 |
| | AUROC | 0.872 ± 0.005 | 0.864 ± 0.005 | 0.828 ± 0.006 | 0.855 ± 0.005 | 0.835 ± 0.004 |
| Test | balanced accuracy | 0.348 ± 0.003 | 0.350 ± 0.003 | 0.356 ± 0.004 | 0.395 ± 0.005 | 0.464 ± 0.009 |
| | precision | 0.321 ± 0.005 | 0.314 ± 0.004 | 0.258 ± 0.003 | 0.250 ± 0.004 | 0.213 ± 0.002 |
| | recall | 0.348 ± 0.003 | 0.350 ± 0.003 | 0.356 ± 0.004 | 0.395 ± 0.005 | 0.464 ± 0.009 |
| | F1 score | 0.209 ± 0.003 | 0.197 ± 0.004 | 0.191 ± 0.004 | 0.205 ± 0.005 | 0.203 ± 0.003 |
| | AUROC | 0.864 ± 0.006 | 0.855 ± 0.006 | 0.827 ± 0.007 | 0.852 ± 0.006 | 0.815 ± 0.007 |

Table S19: Off-time kinetics classification head performance for the ablated model without the dynamic contact module (DCM) (macro-averaged metrics). Values are reported as mean ± standard error.

| Dataset | Metric | 320 K | 348 K | 379 K | 413 K | 450 K |
|---|---|---|---|---|---|---|
| Training | balanced accuracy | 0.419 ± 0.001 | 0.415 ± 0.001 | 0.423 ± 0.001 | 0.411 ± 0.001 | 0.433 ± 0.001 |
| | precision | 0.380 ± 0.001 | 0.382 ± 0.001 | 0.372 ± 0.001 | 0.336 ± 0.001 | 0.326 ± 0.001 |
| | recall | 0.419 ± 0.001 | 0.415 ± 0.001 | 0.423 ± 0.001 | 0.411 ± 0.001 | 0.433 ± 0.001 |
| | F1 score | 0.359 ± 0.001 | 0.348 ± 0.001 | 0.350 ± 0.001 | 0.313 ± 0.001 | 0.319 ± 0.001 |
| | AUROC | 0.828 ± 0.001 | 0.835 ± 0.001 | 0.804 ± 0.001 | 0.792 ± 0.001 | 0.761 ± 0.001 |
| Validation | balanced accuracy | 0.419 ± 0.003 | 0.416 ± 0.003 | 0.423 ± 0.003 | 0.416 ± 0.003 | 0.440 ± 0.004 |
| | precision | 0.375 ± 0.004 | 0.378 ± 0.004 | 0.370 ± 0.004 | 0.340 ± 0.003 | 0.329 ± 0.003 |
| | recall | 0.419 ± 0.003 | 0.416 ± 0.003 | 0.423 ± 0.003 | 0.416 ± 0.003 | 0.440 ± 0.004 |
| | F1 score | 0.354 ± 0.005 | 0.345 ± 0.005 | 0.346 ± 0.005 | 0.316 ± 0.004 | 0.325 ± 0.003 |
| | AUROC | 0.830 ± 0.005 | 0.838 ± 0.005 | 0.808 ± 0.004 | 0.793 ± 0.005 | 0.765 ± 0.005 |
| Test | balanced accuracy | 0.416 ± 0.004 | 0.409 ± 0.004 | 0.418 ± 0.004 | 0.407 ± 0.004 | 0.429 ± 0.004 |
| | precision | 0.373 ± 0.004 | 0.373 ± 0.005 | 0.364 ± 0.005 | 0.332 ± 0.003 | 0.325 ± 0.003 |
| | recall | 0.416 ± 0.004 | 0.409 ± 0.004 | 0.418 ± 0.004 | 0.407 ± 0.004 | 0.429 ± 0.004 |
| | F1 score | 0.350 ± 0.005 | 0.339 ± 0.006 | 0.340 ± 0.006 | 0.307 ± 0.005 | 0.319 ± 0.003 |
| | AUROC | 0.820 ± 0.006 | 0.827 ± 0.006 | 0.800 ± 0.005 | 0.785 ± 0.006 | 0.754 ± 0.006 |

Table S20: Performance comparison on the mdCATH test set at 320 K. Dynamic contact classification metrics are reported for all models, along with RMSE for frequency (contact occupancy) prediction. AUROC is only reported for ESMDynamic, as probabilistic outputs are not available for the other models. Values are shown as mean ± standard error. Best performance for each metric is shown in bold.

| Metric | ESMDynamic | BioEmu | AlphaFlow | ESMFlow (base) | ESMFlow (distilled) |
|---|---|---|---|---|---|
| Balanced Acc. | **0.796 ± 0.007** | 0.794 ± 0.006 | 0.687 ± 0.005 | 0.670 ± 0.005 | 0.593 ± 0.003 |
| Precision | 0.511 ± 0.012 | 0.568 ± 0.011 | **0.839 ± 0.011** | 0.764 ± 0.015 | 0.789 ± 0.016 |
| Recall | **0.767 ± 0.010** | 0.739 ± 0.010 | 0.402 ± 0.010 | 0.408 ± 0.012 | 0.248 ± 0.011 |
| F1 | 0.569 ± 0.008 | **0.598 ± 0.009** | 0.498 ± 0.008 | 0.453 ± 0.009 | 0.296 ± 0.008 |
| AUROC | 0.889 ± 0.006 | – | – | – | – |
| RMSE | 0.076 ± 0.002 | **0.063 ± 0.002** | 0.071 ± 0.002 | 0.084 ± 0.003 | 0.099 ± 0.003 |

Table S21: Performance comparison on the ATLAS test set. Dynamic contact classification metrics are reported for all models, along with RMSE for frequency (contact occupancy) prediction. AUROC is only reported for ESMDynamic, as probabilistic outputs are not available for the other models. Values are shown as mean ± standard error.

| Metric | ESMDynamic | BioEmu | AlphaFlow | ESMFlow (base) | ESMFlow (distilled) |
|---|---|---|---|---|---|
| Balanced Acc. | **0.872 ± 0.010** | 0.770 ± 0.011 | 0.780 ± 0.006 | 0.749 ± 0.007 | 0.685 ± 0.007 |
| Precision | 0.284 ± 0.013 | 0.450 ± 0.022 | **0.675 ± 0.020** | 0.552 ± 0.029 | 0.548 ± 0.034 |
| Recall | **0.890 ± 0.011** | 0.664 ± 0.028 | 0.582 ± 0.013 | 0.557 ± 0.015 | 0.447 ± 0.019 |
| F1 | 0.413 ± 0.016 | 0.458 ± 0.015 | **0.597 ± 0.011** | 0.490 ± 0.018 | 0.387 ± 0.018 |
| AUROC | 0.942 ± 0.007 | – | – | – | – |
| RMSE | 0.063 ± 0.003 | 0.050 ± 0.003 | **0.044 ± 0.003** | 0.058 ± 0.004 | 0.072 ± 0.004 |

Table S22: Comparison of ESMFold-predicted native contacts and ground-truth native contacts from the PDB as baselines for dynamic contact prediction across mdCATH temperature conditions. Metrics are reported as mean ± standard error.

| T (K) | Metric | ESMFold | Native contacts (PDB) |
|---|---|---|---|
| 320 | Balanced Acc. | 0.609 ± 0.003 | **0.627 ± 0.001** |
| | Precision | 0.639 ± 0.011 | **0.646 ± 0.002** |
| | Recall | 0.238 ± 0.007 | **0.273 ± 0.001** |
| | F1 | 0.320 ± 0.007 | **0.358 ± 0.001** |
| 348 | Balanced Acc. | 0.591 ± 0.003 | **0.604 ± 0.001** |
| | Precision | 0.720 ± 0.011 | **0.727 ± 0.002** |
| | Recall | 0.197 ± 0.006 | **0.224 ± 0.001** |
| | F1 | 0.288 ± 0.007 | **0.321 ± 0.001** |
| 379 | Balanced Acc. | 0.568 ± 0.003 | **0.577 ± 0.001** |
| | Precision | **0.821 ± 0.009** | 0.820 ± 0.002 |
| | Recall | 0.150 ± 0.005 | **0.169 ± 0.001** |
| | F1 | 0.238 ± 0.006 | **0.265 ± 0.001** |
| 413 | Balanced Acc. | 0.534 ± 0.005 | **0.537 ± 0.001** |
| | Precision | 0.909 ± 0.007 | **0.914 ± 0.001** |
| | Recall | 0.101 ± 0.004 | **0.112 ± 0.001** |
| | F1 | 0.174 ± 0.005 | **0.191 ± 0.001** |
| 450 | Balanced Acc. | **0.498 ± 0.007** | 0.492 ± 0.002 |
| | Precision | 0.976 ± 0.004 | **0.978 ± 0.001** |
| | Recall | 0.062 ± 0.002 | **0.068 ± 0.0005** |
| | F1 | 0.115 ± 0.003 | **0.124 ± 0.001** |

Table S23: Performance comparison between ESMDynamic and BioEmu across BioEmu benchmark datasets. Values are reported as mean ± standard deviation. Best performance for each metric is shown in bold. The set "cryptic pocket (res. range)" only considers dynamic contacts involving at least one residue from the pocket.

| Benchmark | Model | Balanced Acc. | Precision | Recall | F1 |
|---|---|---|---|---|---|
| OOD60 | ESMDynamic | **0.854 ± 0.079** | 0.024 ± 0.018 | **0.898 ± 0.148** | 0.046 ± 0.034 |
| | BioEmu | 0.795 ± 0.127 | **0.062 ± 0.051** | 0.775 ± 0.264 | **0.106 ± 0.081** |
| Domain motion | ESMDynamic | **0.903 ± 0.104** | 0.015 ± 0.013 | **0.879 ± 0.212** | 0.029 ± 0.025 |
| | BioEmu | 0.838 ± 0.147 | **0.048 ± 0.036** | 0.697 ± 0.303 | **0.083 ± 0.054** |
| Cryptic pocket | ESMDynamic | **0.921 ± 0.051** | 0.016 ± 0.014 | **0.912 ± 0.096** | 0.031 ± 0.027 |
| | BioEmu | 0.913 ± 0.087 | **0.037 ± 0.024** | 0.857 ± 0.178 | **0.068 ± 0.041** |
| Cryptic pocket (res. range) | ESMDynamic | **0.926 ± 0.057** | 0.032 ± 0.026 | **0.889 ± 0.186** | 0.061 ± 0.047 |
| | BioEmu | 0.918 ± 0.098 | **0.060 ± 0.036** | 0.843 ± 0.250 | **0.109 ± 0.061** |
| Local unfolding | ESMDynamic | – | – | **0.940 ± 0.090** | – |
| | BioEmu | – | – | 0.931 ± 0.141 | – |

#### 4 Supplementary Figures

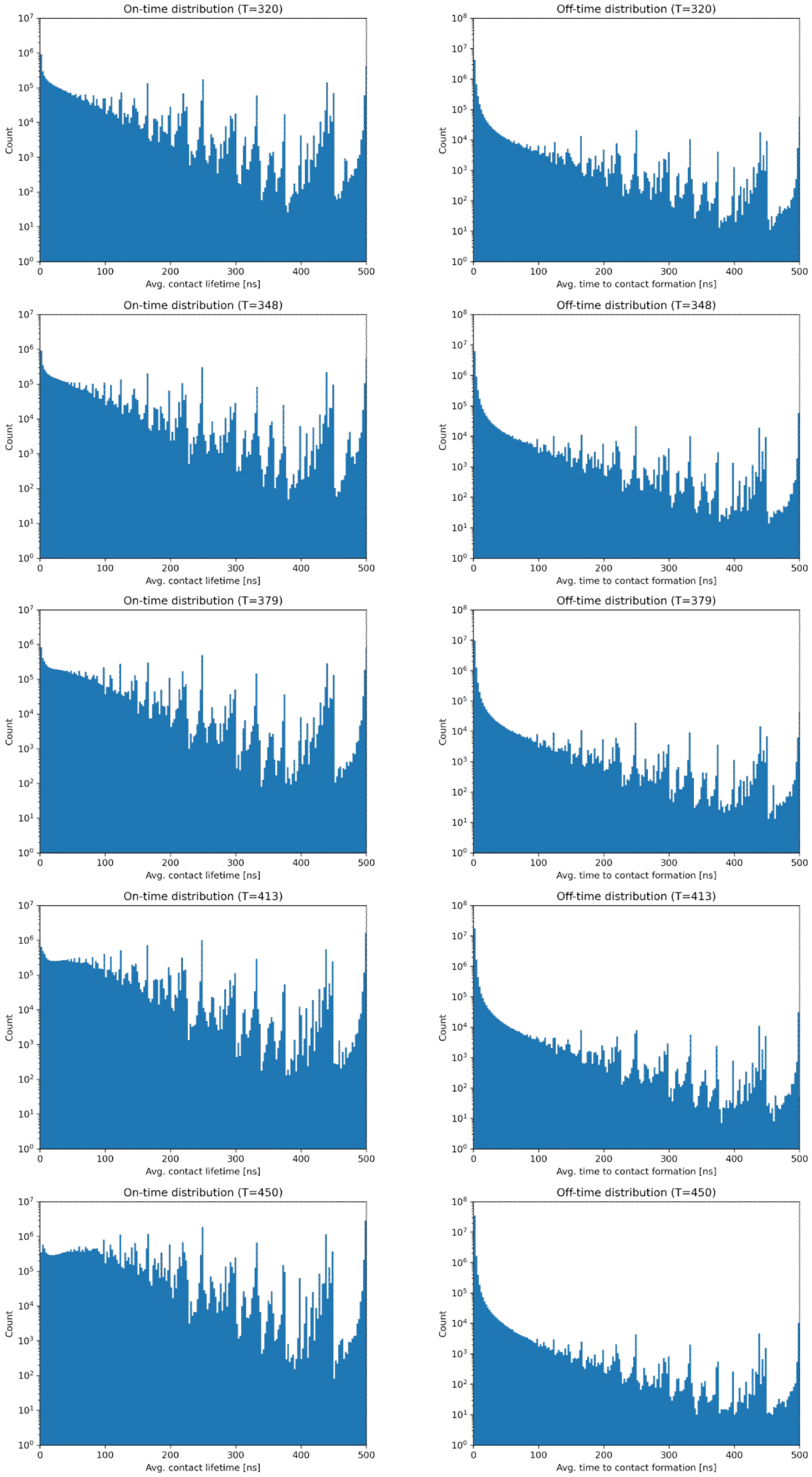


Figure S1: Distribution of on-time (contact lifetime, left) and off-time (time to formation, right) for dynamic contacts in mdCATH dataset across temperatures (rows).

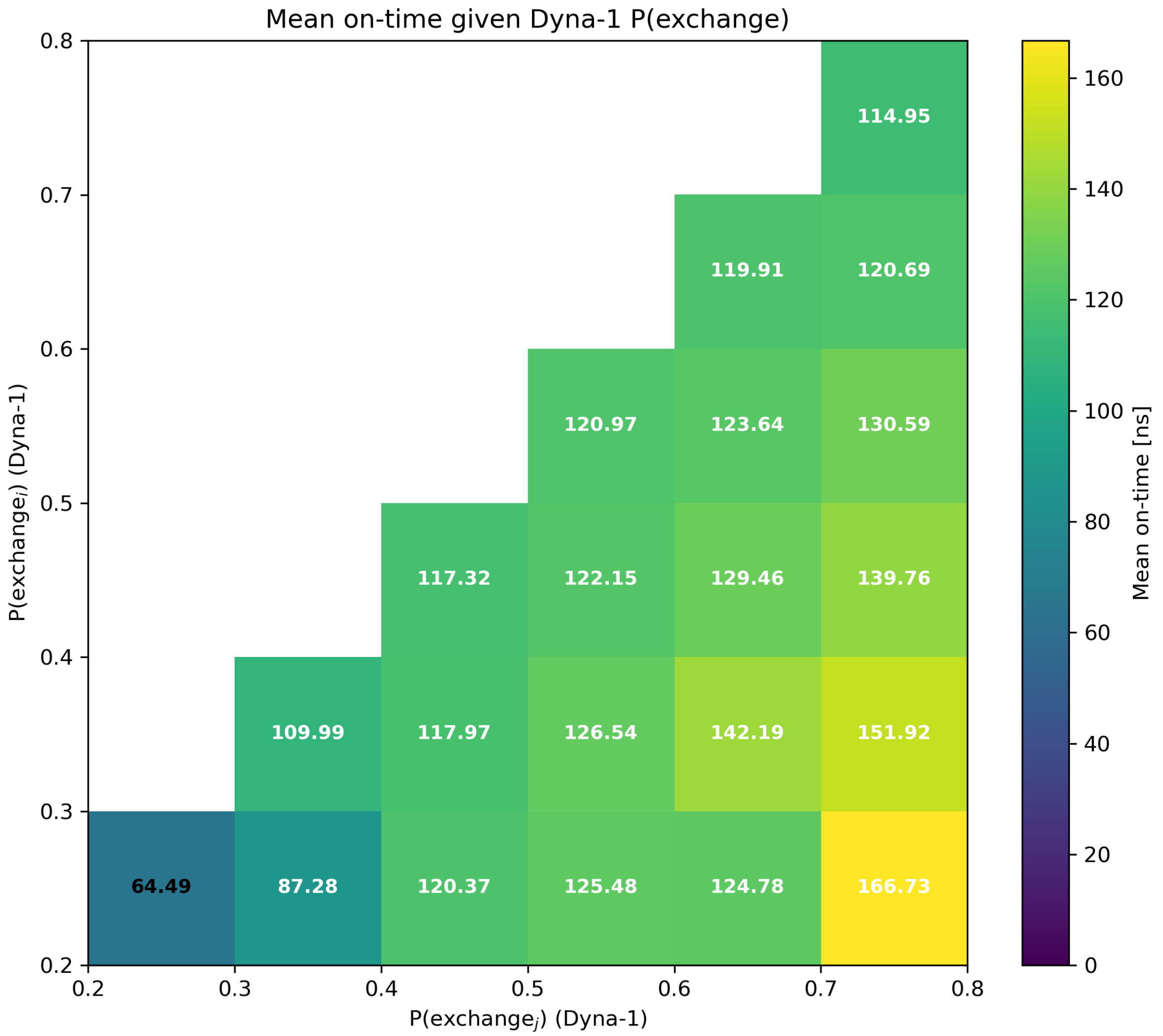


Figure S2: Relationship between residue-level dynamics inferred by Dyna-1 and contact lifetimes in mdCATH (320 K). Each bin corresponds to residue pairs grouped by the Dyna-1 scores of residues $i$ and $j$, with values indicating the average time a contact remains formed (on-time). Contacts involving at least one residue with a high Dyna-1 score tend to exhibit longer lifetimes within the accessible MD timescales.

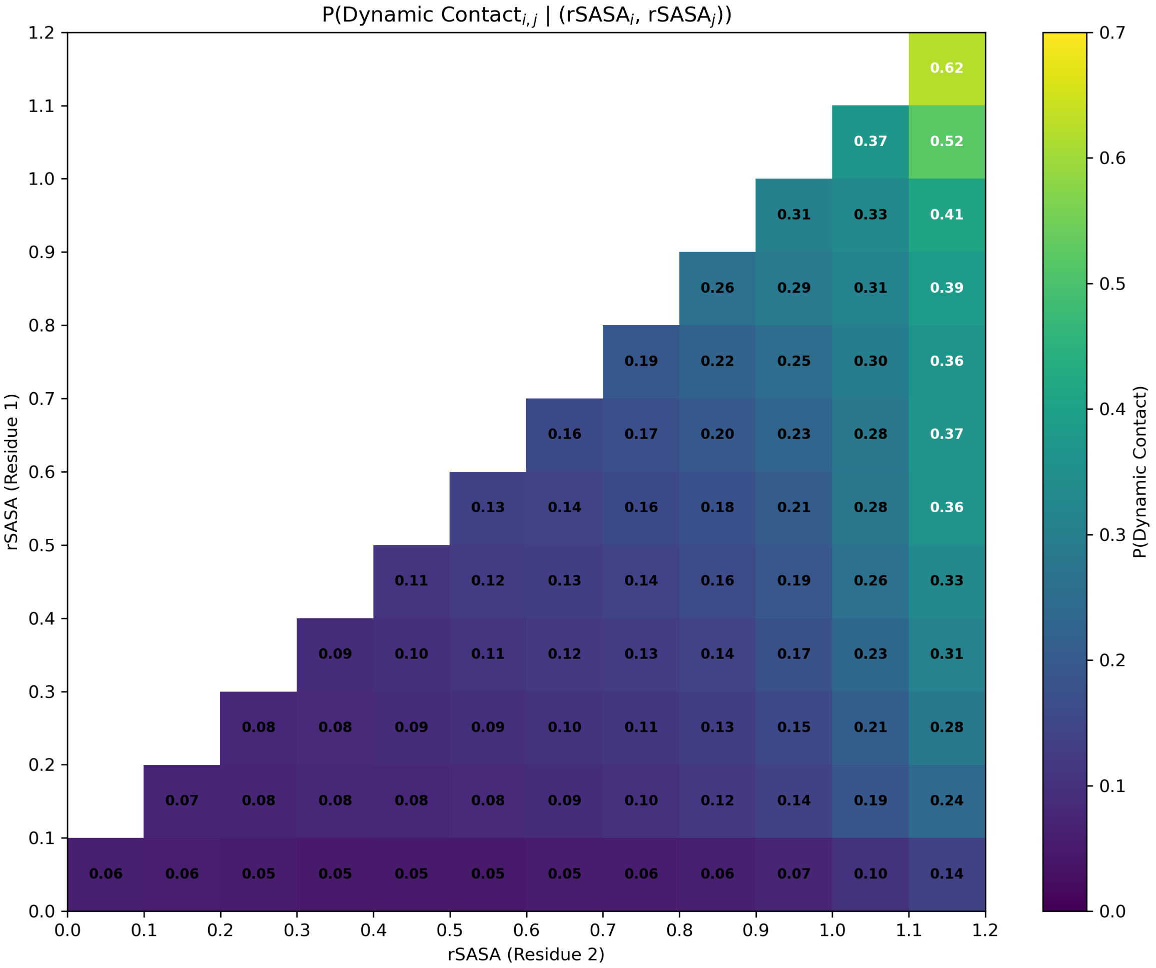


Figure S3: Conditional probability of dynamic contact formation as a function of relative residue solvent accessibility (rSASA). Dynamic-contact probability is shown as a function of the rSASA of residue pair $(i, j)$.

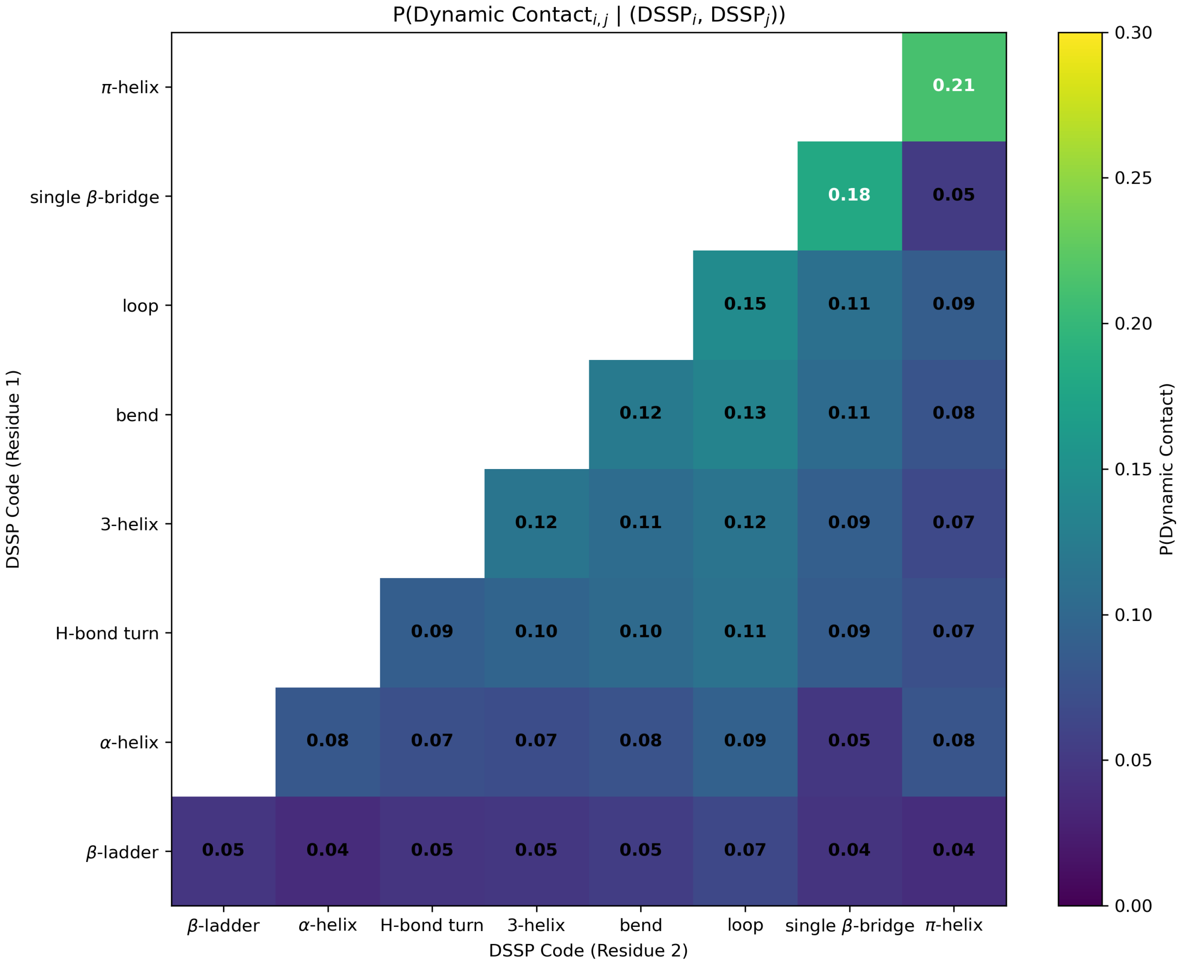


Figure S4: Conditional probability of dynamic contact formation as a function of secondary structure. Dynamic-contact probability is shown as a function of the DSSP assignments of residue pair $(i, j)$.

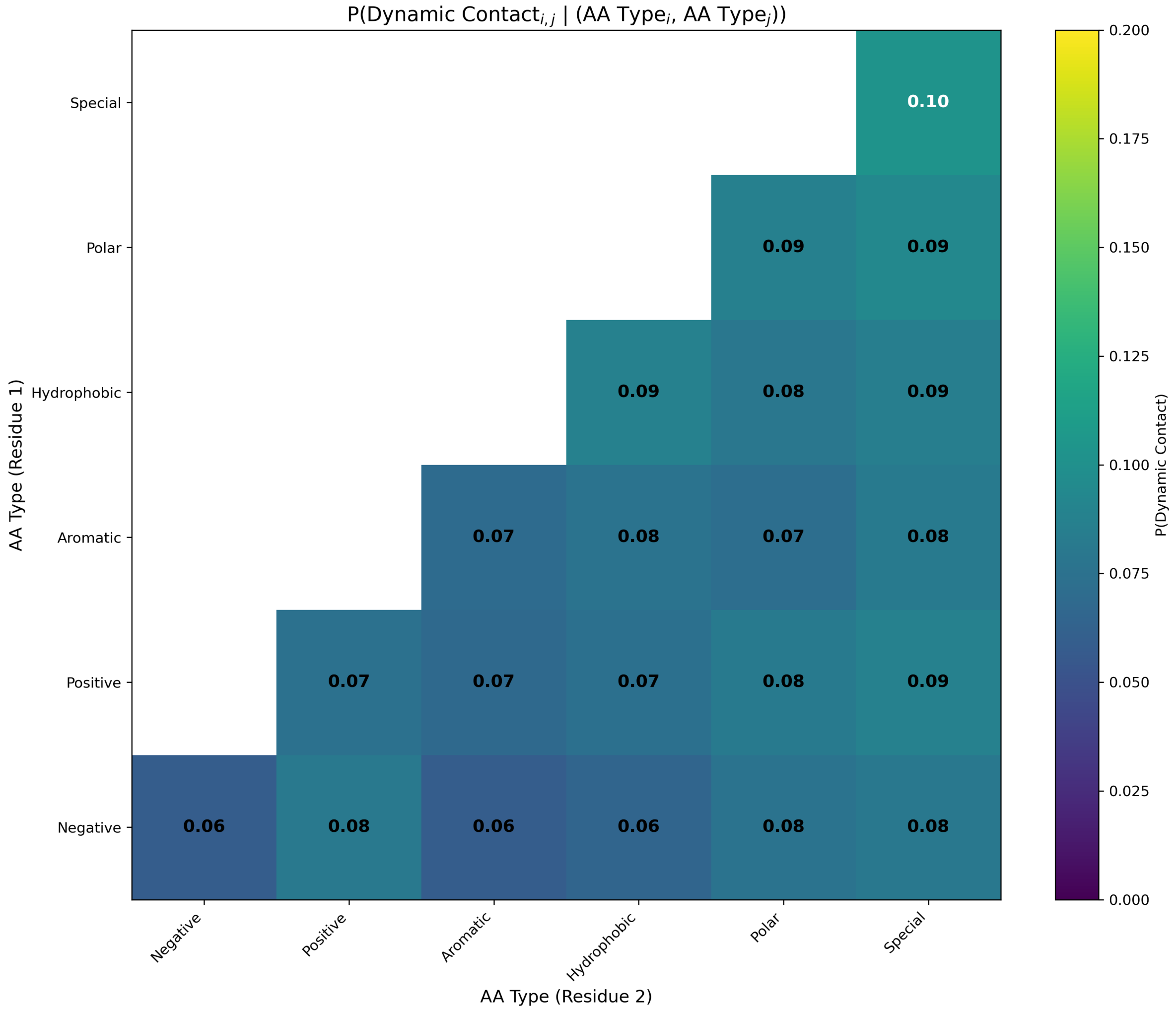


Figure S5: Conditional probability of dynamic contact formation as a function of amino acid type. Dynamic-contact probability is shown as a function of the amino acid types of residue pair $(i,\ j)$.

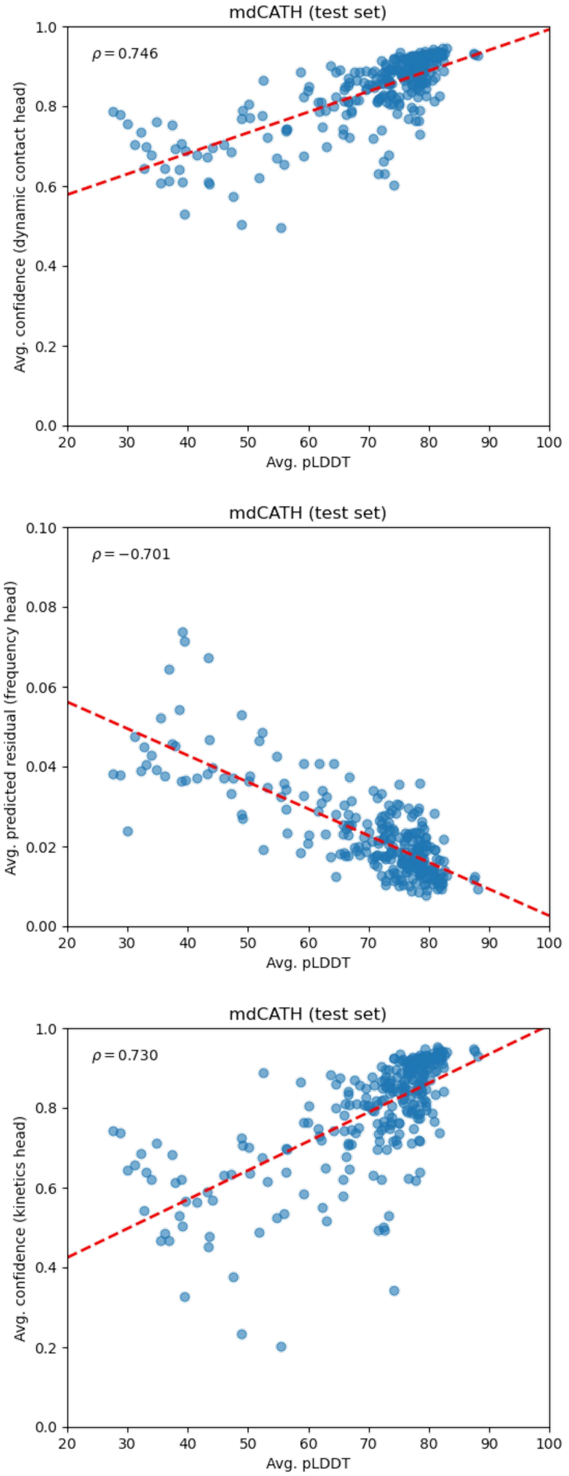


Figure S6: Correlation between average predicted confidence or error (ESMDynamic) and average pLDDT (ESMFold). Spearman's $\rho$ is shown as the correlation metric; best linear fit displayed as visual aid.

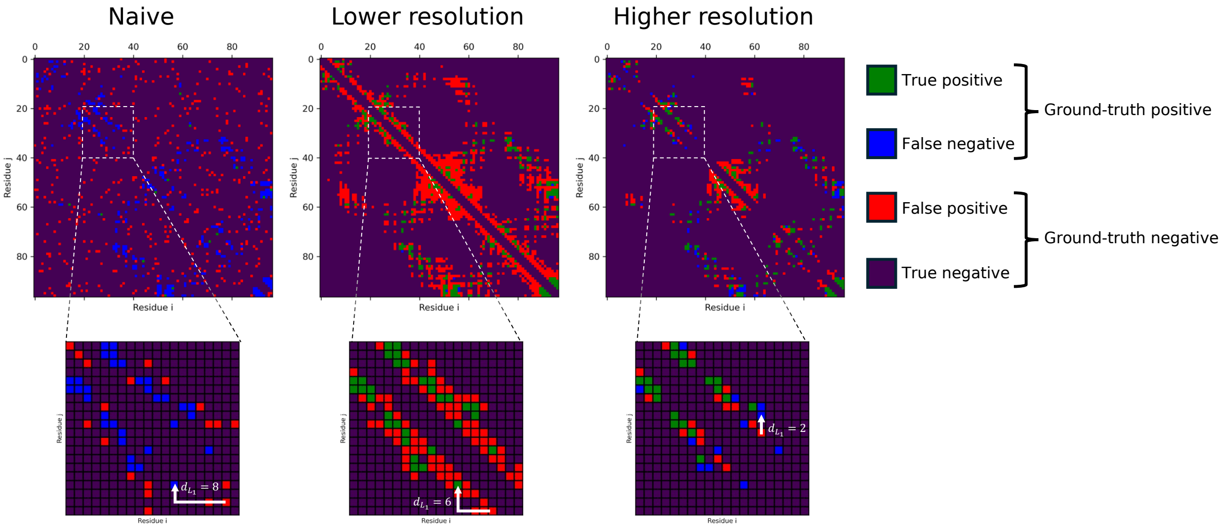


Figure S7: Illustrative example of minimum $L_1$ distance computation in sequence space. (Left) A random naive model yields many false positives, resulting in a low resolution score. (Center) A lower-resolution model achieves higher recall but lower precision; many false positives are predicted, some far from ground-truth positives. (Right) A higher-resolution model predicts fewer false positives, improving precision at the cost of recall, with some false negatives. Predicted positives are more tightly clustered around the ground-truth.

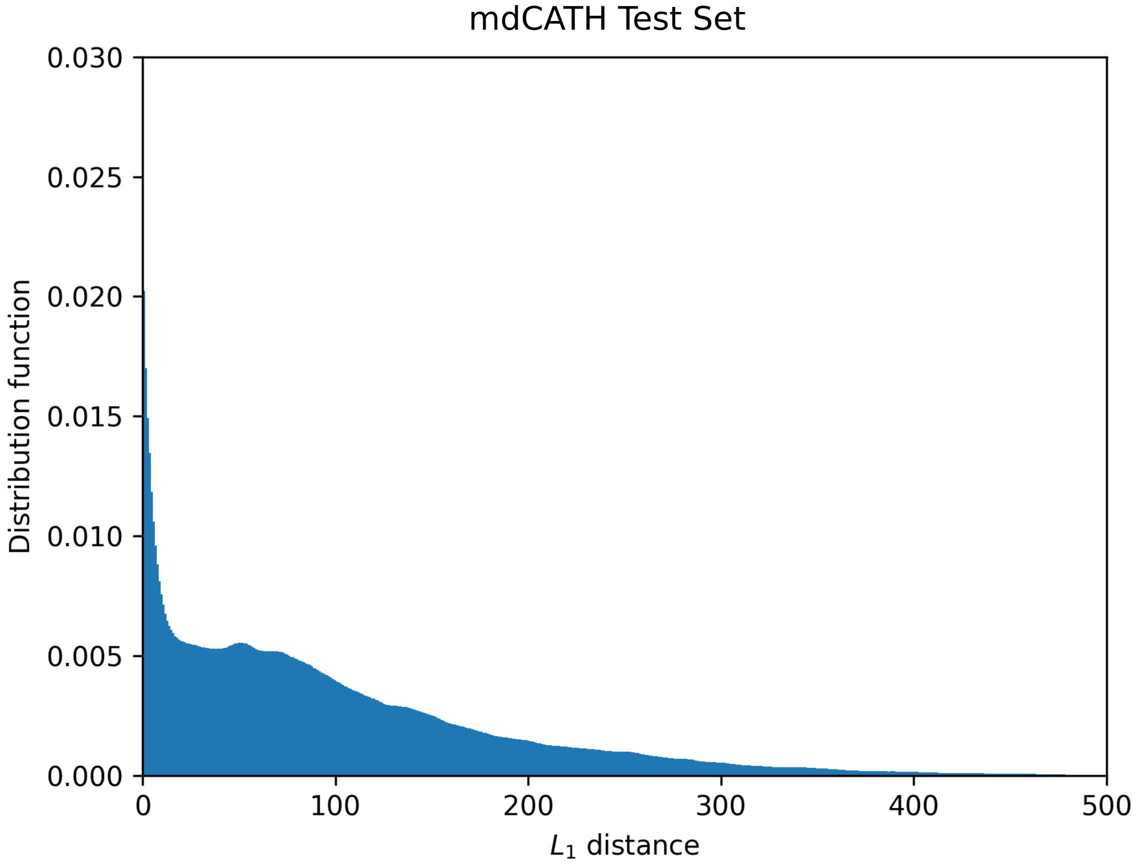


Figure S8: $L_1$ radial distribution function for dynamic contacts in the mdCATH test set. Distribution is computed as $f(r) = \frac{1}{N}\langle p(r)/T(r)\rangle$, where $p(r)$ is the number of dynamic contacts at distance $r$ of another dynamic contact, $T(r)$ is the total number of possible residue-residue pairs at the same distance, and $1/N$ normalizes the distribution. The concentration of probability towards lower distances indicates that dynamic contacts tend to cluster together.

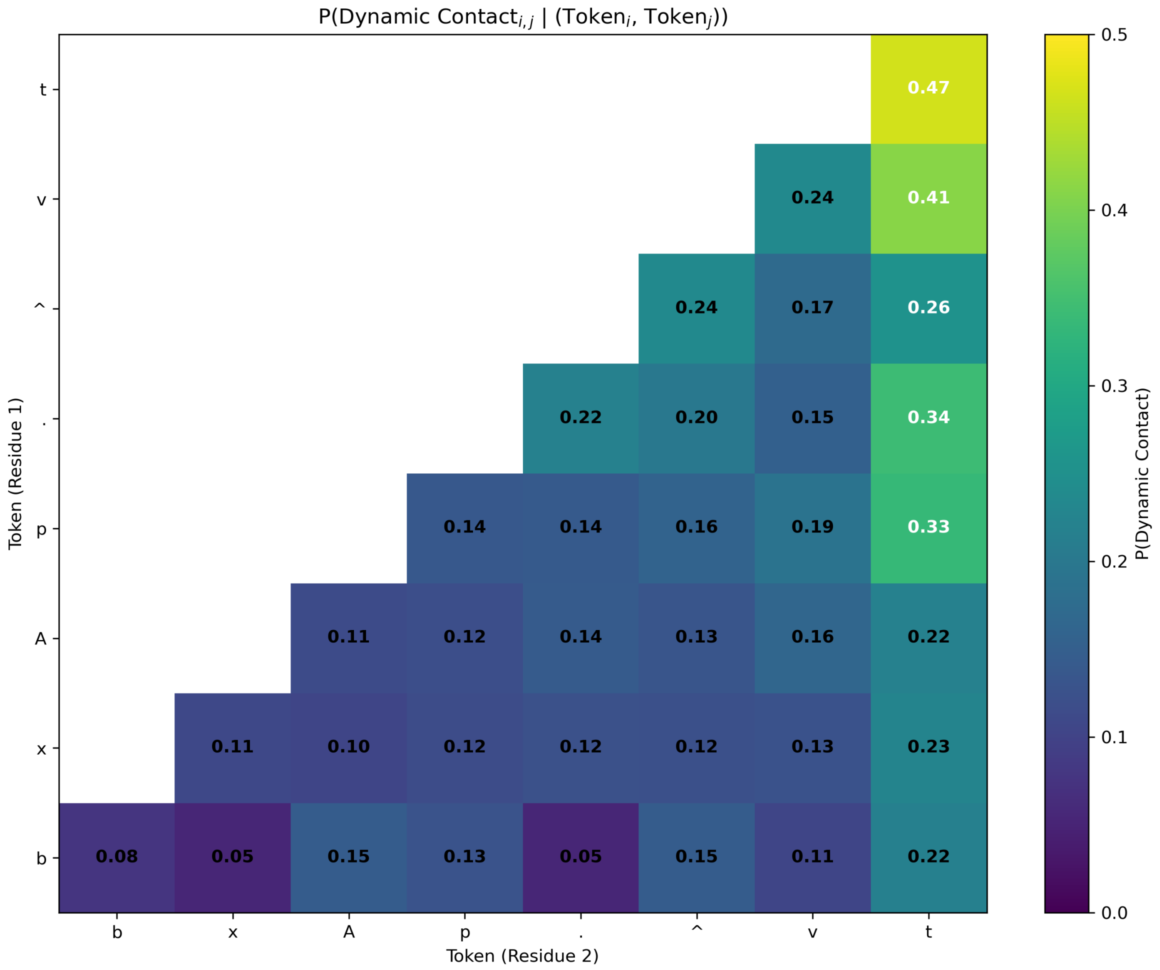


Figure S9: Conditional probability of dynamic contact prediction as a function of residue-level dynamics annotations (RelaxDB tokens). Dynamic contact prediction probability is shown as a function of the RelaxDB tokens assigned to residue pair $(i, j)$. Tokens correspond to distinct dynamical regimes: $A$ (no detectable dynamics), $v$ (fast ps–ns motion), ^ (slow $\mu$s–ms conformational exchange), $b$ (combined fast and slow motion), $t$ (disordered terminus), $p$ (proline), $x$ (no reported data), and . (missing assignment). Elevated dynamic contact probabilities are observed for residues associated with flexibility or disorder, particularly those displaying terminal disorder, fast ps–ns motions, or slower conformational exchange.

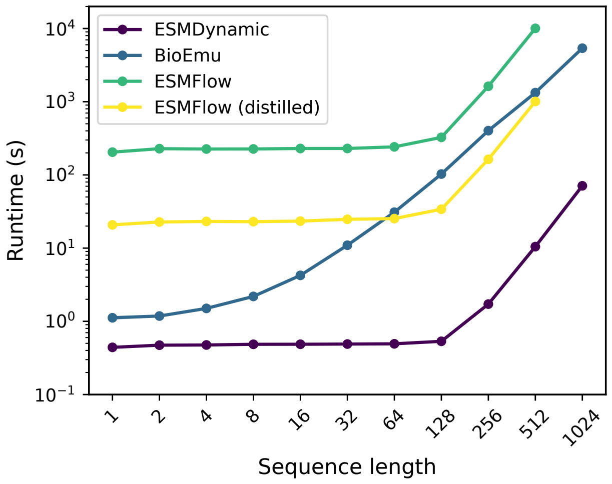


Figure S10: Inference wall-clock time as a function of protein sequence length (1–1024 residues) for ESMDynamic and representative generative models (BioEmu, ESMFlow, and distilled ESMFlow). ESMDynamic exhibits consistently lower runtime across all sequence lengths, requiring only a single forward pass per sequence. In contrast, generative approaches incur substantially higher computational cost due to repeated sampling (250 samples per protein here), resulting in longer runtimes. Chunk size of 256 was used for ESMDynamic. Values are averages of 5 replicates.

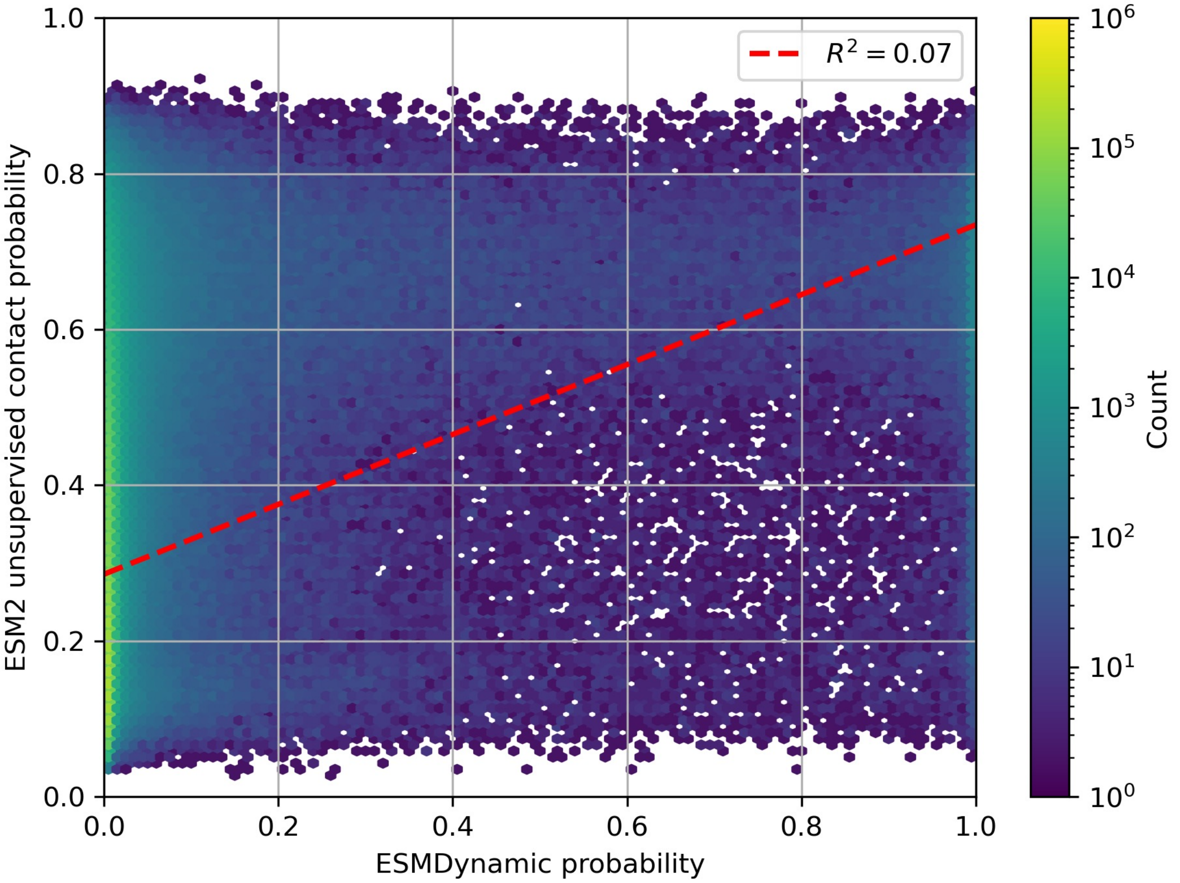


Figure S11: ESMDynamic's probability outputs show very weak correlation with ESM2's unsupervised contact probabilities, indicating that the model captures novel information related to protein dynamics rather than simply reproducing native contact probabilities.

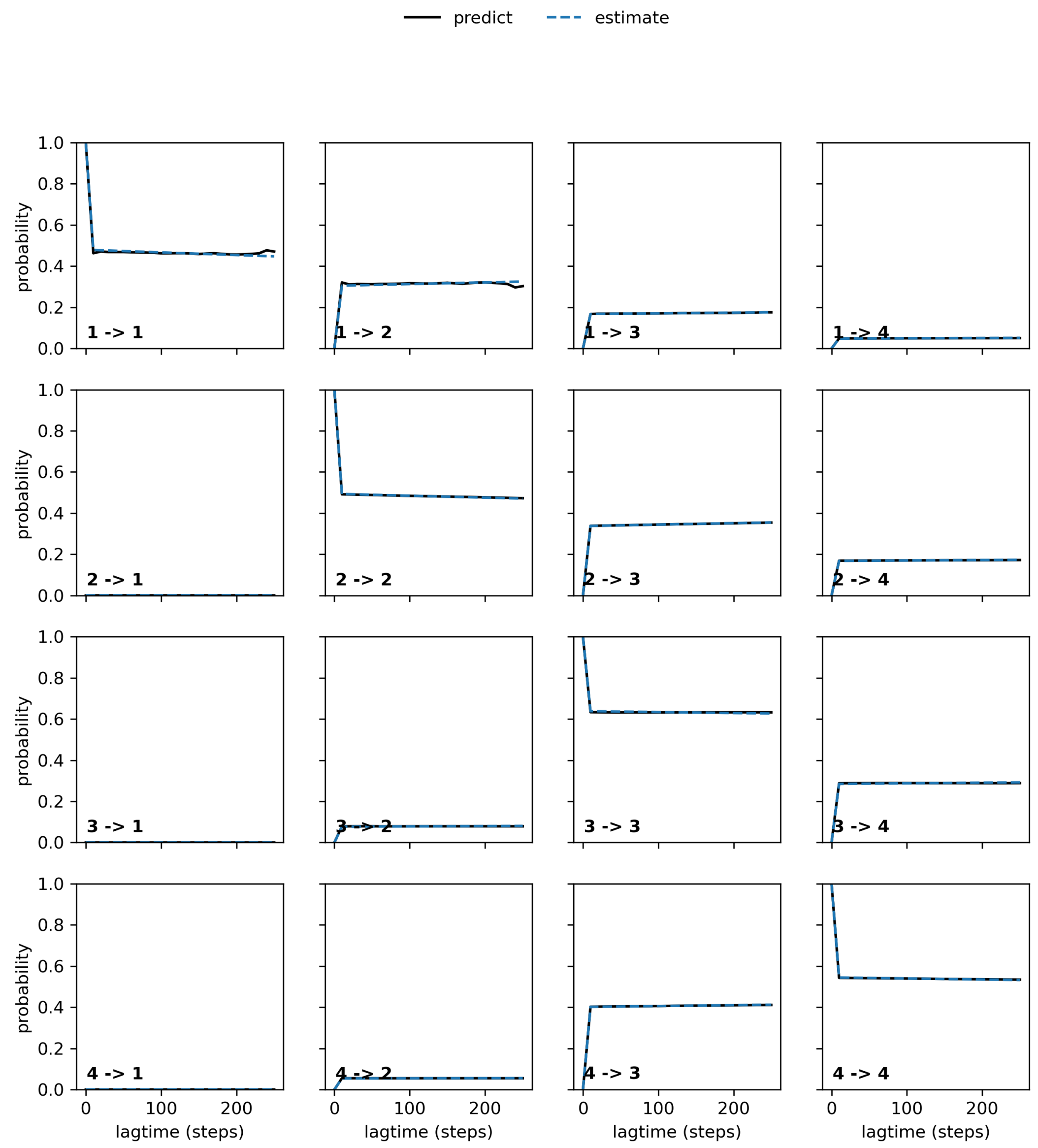


Figure S12: Chapman-Kolmogorov test for OsSWEET2b MSM with four macrostates. Agreement between estimated and predicted transition probabilities at increasing lagtime indicate that the fitted MSM obeys the Markovian property.

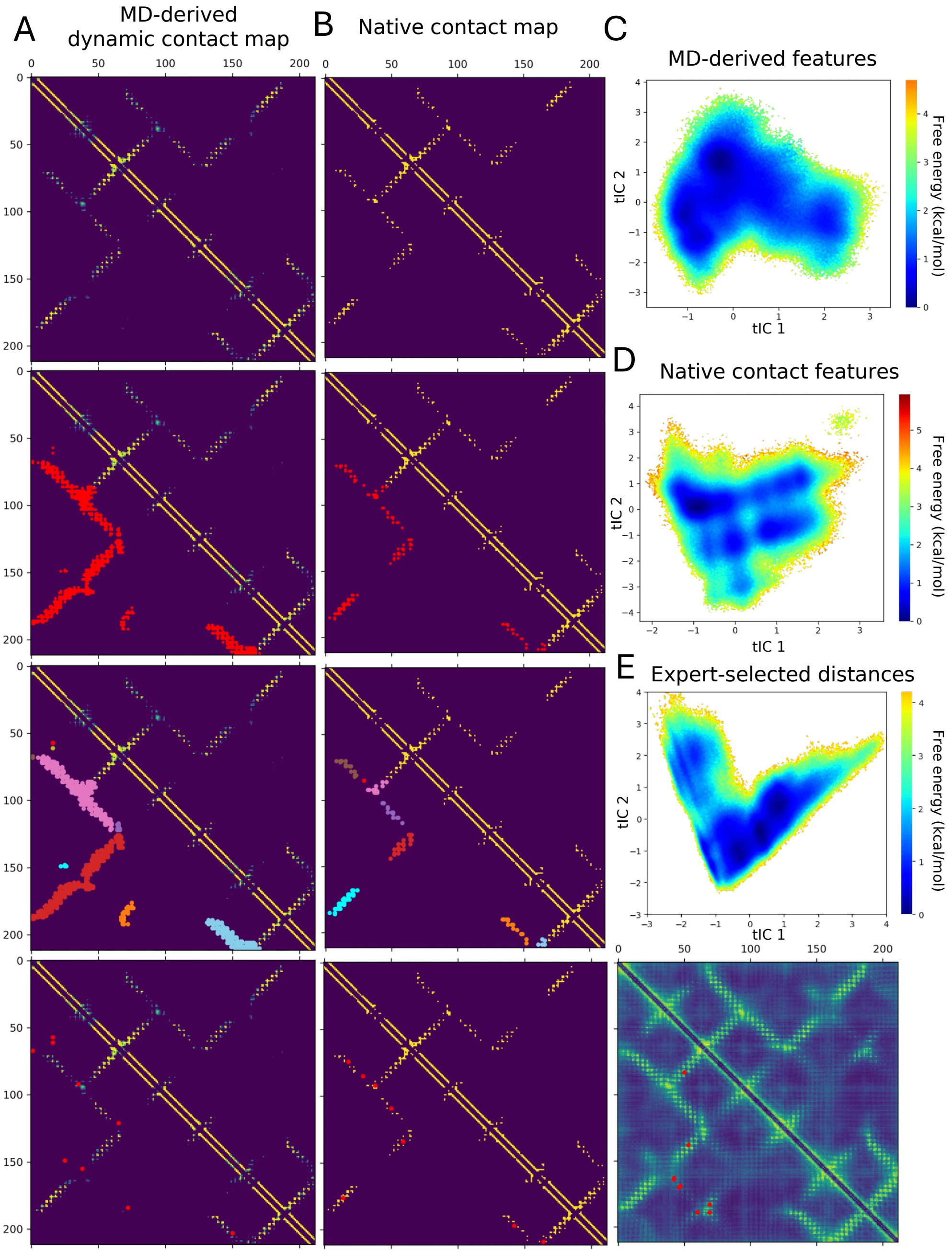


Figure S13: Alternative feature selection strategies using MD-derived and native contact baselines. (A–B) Collective variable (CV) selection pipeline applied to (A) MD-derived dynamic contact maps and (B) native contact maps. In both cases, the selected CVs differ substantially from expert-curated features and those obtained from ESMDynamic. (C–D) Free energy landscapes constructed from the CVs identified in (A) and (B), respectively. Both landscapes deviate from those obtained using expert-selected or ESMDynamic-derived CVs, indicating that these baseline feature sets fail to resolve the underlying transport mechanism. (E) Top: Free energy landscape derived from expert-selected CVs projected on top two tICs, showing qualitative agreement with the ESMDynamic-based landscape. Bottom: corresponding expert-selected CVs for reference.

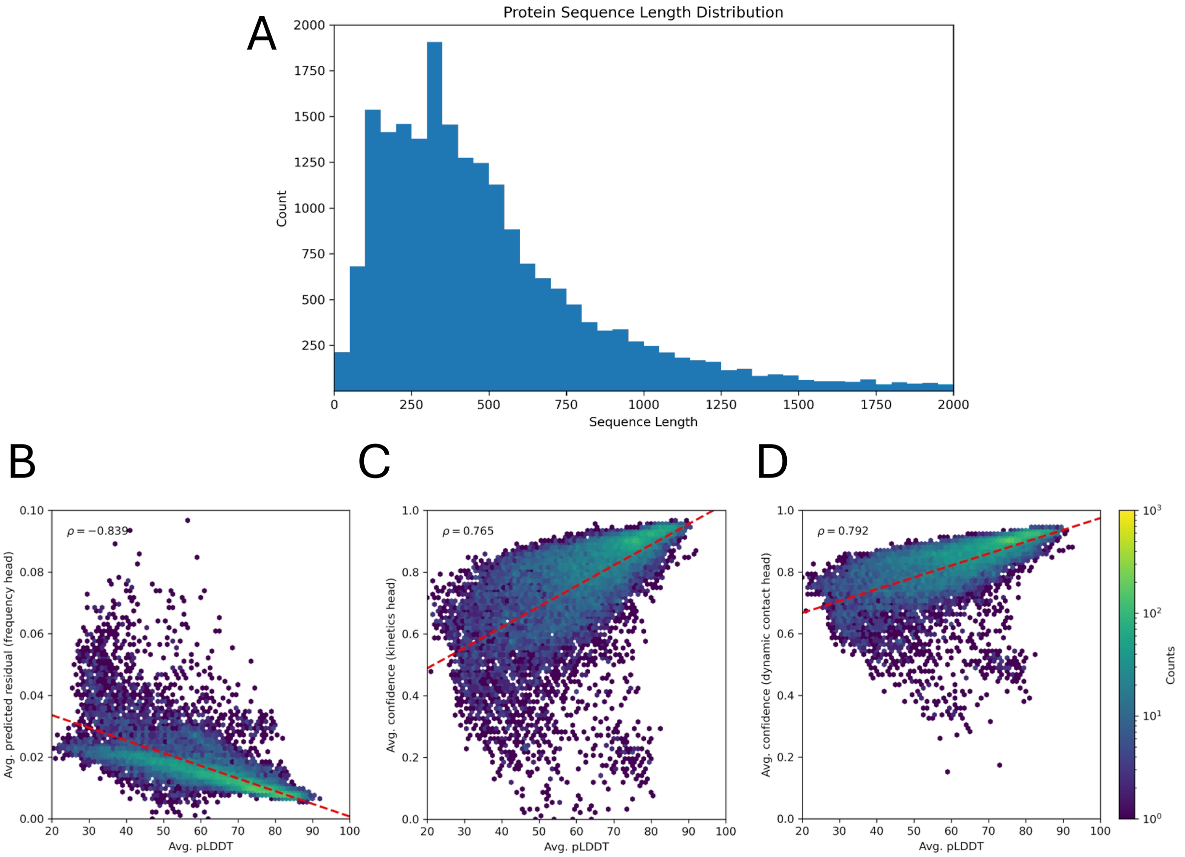


Figure S14: Large-scale application of ESMDynamic to the human proteome (UniProt UP000005640). (A) Distribution of protein sequence lengths, showing that the majority of proteins fall below the 1000-residue cutoff used for inference (covering ∼88% of the proteome). (B–D) Relationship between ESMDynamic uncertainty outputs and ESMFold confidence (pLDDT). Spearman's $\rho$ is shown as the correlation metric; best linear fit displayed as visual aid. (B) Predicted error of the frequency (equilibrium probability) head shows negative correlation with pLDDT, indicating that high confidence structures lead to lower estimated error. (C) Confidence scores from the kinetics head increase with pLDDT. (D) Confidence scores from the dynamic contact classification head show a similar positive correlation with pLDDT.

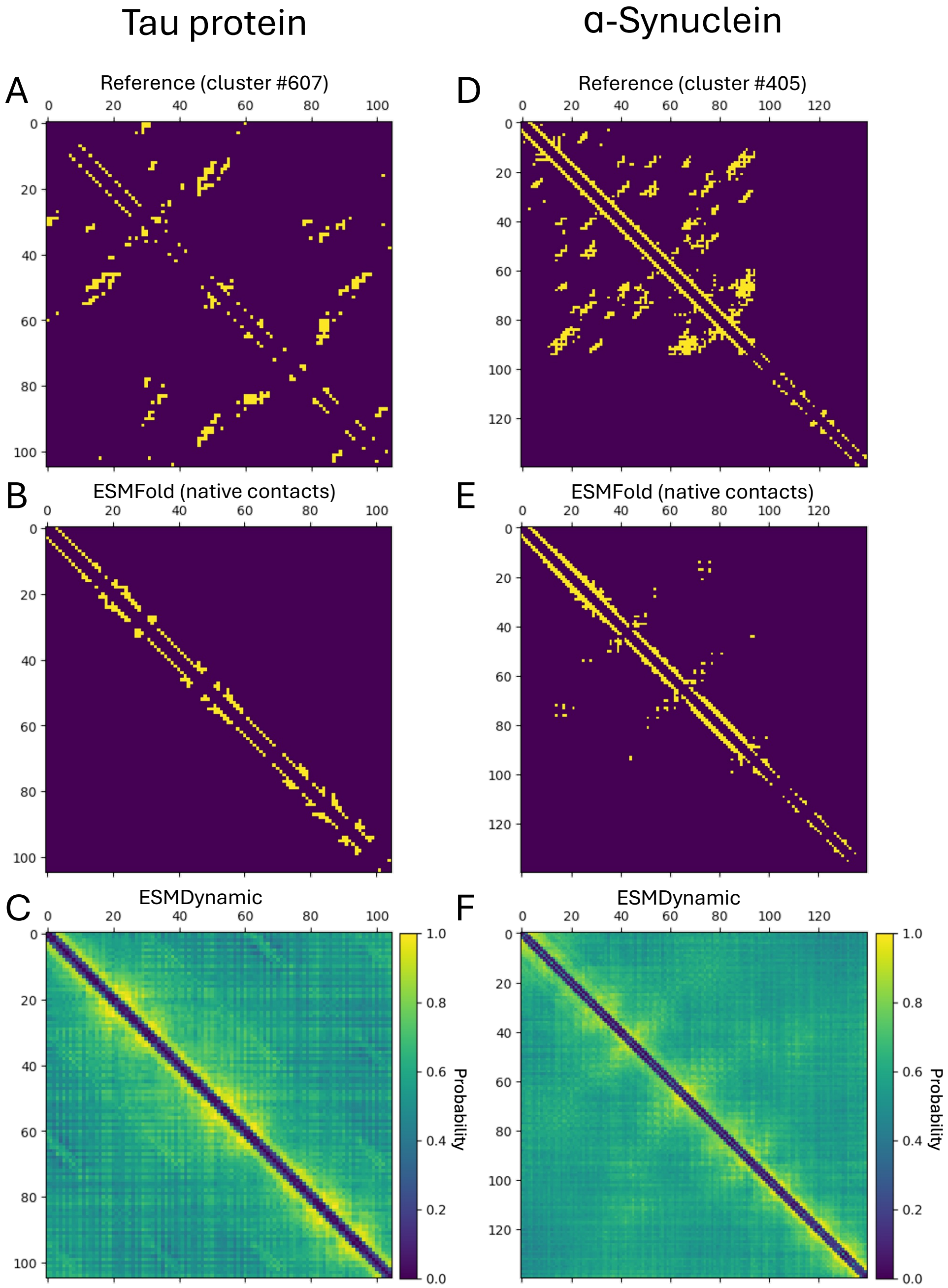


Figure S15: ESMDynamic results on intrinsically disordered proteins. (A–C) Tau protein. (A) Dynamic contact map derived from experimental structures. (B) Native contact map from ESMFold. (C) Dynamic contact map from ESMDynamic. (D–F) $\alpha$-Synuclein. (D) Dynamic contact map derived from experimental structures. (E) Native contact map from ESMFold. (F) Dynamic contact map from ESMDynamic.

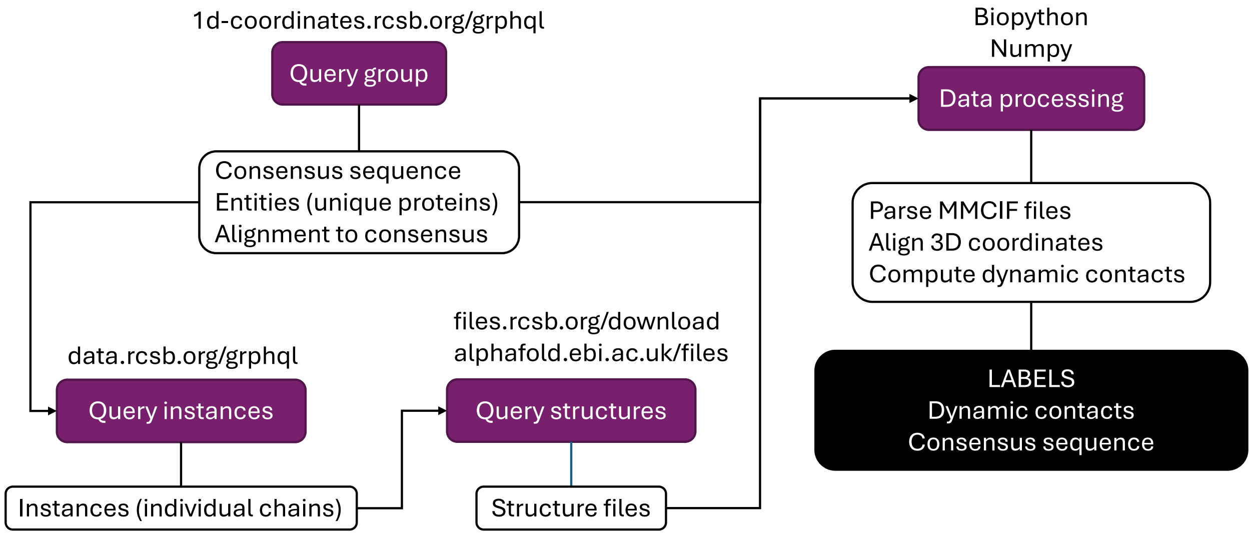


Figure S16: Workflow for obtaining structural clusters from the RCSB PDB. An entity refers to a unique resolved structure (i.e., an independent PDB entry), while an instance corresponds to a specific protein chain within that structure (multiple instances can belong to the same entity). First, consensus sequences from sequence clusters (at 95% identity) are retrieved, along with structure identifiers (entity IDs) and alignments of cluster members to the consensus. Next, instance IDs corresponding to each entity are obtained. The 3D coordinates of each chain instance are then downloaded and used to compute dynamic contacts. The consensus sequence is retained as the reference for the cluster.

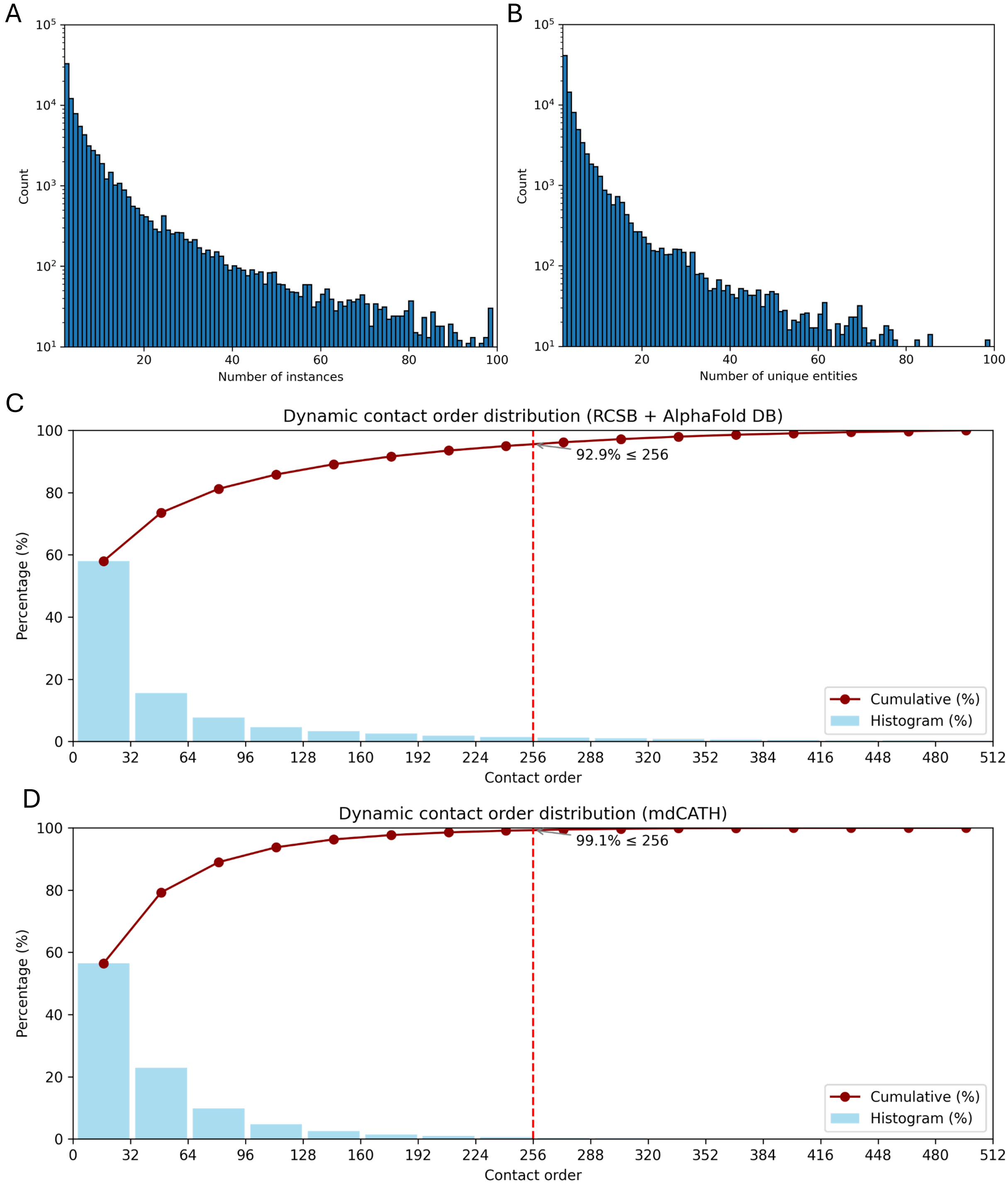


Figure S17: Dataset statistics. (A) Cluster size distribution for RCSB PDB and AlphaFold DB ensembles by number of resolved chains per cluster. (B) Same distribution by number of unique entities (independent structures). (C) Dynamic contact order (sequence distance between contacting residues) for the PDB + AlphaFold DB dataset; 93% occur within 256 residues, supporting this crop size for training. (D) Dynamic contact order for mdCATH, with 99% within 256 residues.
